# Temporal regulation of TAK1 to counteract muscular dystrophy

**DOI:** 10.1101/2022.07.22.501199

**Authors:** Anirban Roy, Tatiana E. Koike, Aniket S. Joshi, Meiricris Tomaz da Silva, Kavya Mathukumalli, Mingfu Wu, Ashok Kumar

**Affiliations:** Department of Pharmacological and Pharmaceutical Sciences, University of Houston College of Pharmacy, Houston, TX 77204, USA

## Abstract

Muscular dystrophy is a group of genetic neuromuscular disorders that involves severe muscle wasting. Transforming growth factor β-activated kinase 1 (TAK1) is an important signaling protein that regulates cell survival, growth, and inflammation. TAK1 has been recently found to promote myofiber growth in skeletal muscle of adult mice. However, the role of TAK1 in muscle disorders remains poorly understood. In the present study, we have investigated how TAK1 affects progression of dystrophic phenotype in the mdx mouse model of Duchnne muscular dystrophy (DMD). TAK1 is highly activated during peak necrotic phase in mdx mice. Targeted inducible inactivation of TAK1 inhibits muscle injury, necroptosis, and accumulation of macrophages in dystrophic muscle of mdx mice. Additionally, targeted inactivation of TAK1 leads to the activation of autophagy and Notch and Wnt signaling in the dystrophic muscle. However, inactivation of TAK1 significantly reduces myofiber size and muscle contractile function in both young and adult mdx mice. Forced activation of TAK1 in skeletal muscle after peak necrotic phase induces myofiber growth and improves muscle histopathology in mdx mice. Our results suggest that targeted activation of TAK1 can ameliorate disease progression and improve muscle growth in DMD.

**One Sentence Summary:** Our results demonstrate that duly regulation of TAK1 activity ameliorates dystrophic phenotype in a mouse model of Duchnne Muscular Dystrophy.

## INTRODUCTION

Duchenne muscular dystrophy (DMD) is a severe neuromuscular disease that occurs due to mutations in DMD gene that encodes the subsarcolemmal protein, dystrophin. Lack of functional dystrophin protein makes sarcolemma vulnerable to contraction-induced injury that results in degeneration and progressive wasting of nearly all muscles and premature death of afflicted individuals *(1-3)*. The DMD pathology onsets with chronic myofiber necrosis and associated inflammatory response that eventually leads to replacement of myofibers by fat and fibrotic tissue *(4)*. While recent advances in gene therapy, such as CRISPR mediated gene editing or by AAV-mediated delivery of mini-dystrophin gene are prospective therapies for DMD, significant obstacles and safety concerns have thwarted translation of these approaches to true therapies *(5-8)*. Current standard-of-care for DMD is focused on reducing inflammation with corticosteroids, which modestly reduces disease progression but has serious side effects *(9)*. Accumulating evidence suggests that strategies controlling autophagy, metabolic regulation, and immune response along with gene therapy can be promising approaches to ameliorate disease progression in DMD patients *(10-14)*.

Destabilization of the dystrophin-glycoprotein complex (DGC) due to dystrophin deficiency inflicts muscle injury and myonecrosis in DMD *(15-17)*. Due to the chronic nature of myofiber degeneration, dystrophic muscle presents a highly complex microenvironment where factors, including immune and other infiltrating cells, inflammatory cytokines, and pro-fibrotic molecules influence myofiber integrity and regeneration *(4)*. Moreover, aberrant activation of proinflammatory signaling pathways, such as nuclear factor-kappa B (NF-κB) and MAPKs, which exacerbates muscle injury and inflammation, is a common feature in dystrophic muscles *(18-23)*. In addition, a number of other signaling pathways, which regulate myofiber regeneration and fibrosis, including Notch, Wnt, and TGF-β signaling are deregulated in skeletal muscle of animal models of DMD *(4, 24-26)*. However, the physiological significance and molecular underpinning that leads to the activation of various signaling pathways in dystrophic muscles remain poorly understood.

TGF-β-activated kinase 1 (TAK1), a member of the MAPK kinase kinase (MAP3K) family, is a central kinase that regulates multiple cellular responses, including cell survival, proliferation, differentiation, inflammation, innate immune response, and development and morphogenesis of various organs *(27, 28)*. Numerous cytokines, growth factors, and microbial products initiate cell signaling which results in the activation of TAK1 signalosome. For activation, TAK1 interacts with adaptor protein TAK1 binding protein (TAB) 1 and either TAB2 or TAB3 and undergo K63-linked ubiquitination that result in a conformational change leading to the autophosphorylation within the activation loop *(29, 30)*. Once activated, TAK1 phosphorylates MKK4 and MKK3/6 that leads to the activation of JNK and p38 MAPK, respectively. Another important phosphorylation target of TAK1 is I kappa B kinase β (IKKβ) which is responsible for the activation of canonical NF-κB signaling pathway *(29, 31)*.

We have previously reported that TAK1 is essential for satellite stem cell survival, self-renewal and myogenic function. Inducible deletion of TAK1 in satellite cells impairs regeneration of skeletal muscle in adult mice *(32)*. Intriguingly, we also found that TAK1 is critical for growth and maintenance of skeletal muscle mass. Mice with germline deletion of *Tak1* in skeletal muscle show perinatal lethality whereas targeted inducible inactivation of TAK1 causes severe muscle wasting in adult mice, which is accompanied with activation of proteolytic pathways, repression in the rate of protein synthesis, redox imbalance, and mitochondrial dysfunction *(33, 34)*. More recently, we demonstrated that TAK1 is essential for integrity of neuromuscular junctions (NMJs) in adult mice. Forced activation of TAK1 in skeletal muscle causes myofiber hypertrophy and also attenuates the loss of muscle mass in response to functional denervation *(35)*. A recent study has demonstrated that TAK1 is activated in skeletal muscle of DMD patients and dystrophic-deficient mdx mice *(36)*. However, the muscle-specific role of TAK1 in disease progression and the mechanisms by which TAK1 regulates dystrophic phenotype in mdx mice remain poorly understood.

Using inducible muscle-specific TAK1-knockout mice, in the present study, we have investigated the role of TAK1 in disease progression in mdx mice. Our results demonstrate that TAK1 is highly activated during necrotic phase and inactivation of TAK1 reduces muscle injury in young mdx mice potentially through inhibiting inflammatory response and necroptosis. However, inactivation of TAK1 strongly inhibits myofiber growth resulting in reduced muscle mass and strength. In contrast, forced activation of TAK1 in dystrophic muscle of mdx mice through intramuscular injection of AAV6 vectors expressing TAK1 and TAB1 stimulates myofiber growth without having any adverse effect on muscle histopathology. Our experiments suggest that duly regulation of TAK1 can be an important therapeutic approach to improve myofiber growth and function in dystrophic muscle.

## RESULTS

### TAK1 is activated in dystrophic muscle of mdx mice

By performing immunoblotting, we first investigated how the levels of TAK1 are changed in the skeletal muscle of dystrophin-deficient mdx mice compared to wild type (wt) mice at different disease status. There was a significant increase in the levels of phosphorylated TAK1 (p-TAK1) and total TAK1 protein in the gastrocnemius (GA) muscle of 4-week-old mdx mice compared to age-matched wild-type mice (**Fig. 1A, B**). Interestingly, there was no significant difference in the levels of p-TAK1 and total TAK1 in GA muscle of 16-week-old wild type and mdx mice (**Fig. 1A, C**). In the mdx mice, skeletal muscle degeneration starts at the age of 2.5 weeks and severe myonecrosis with inflammation is observed around 3.5 to 4 weeks. Thereafter, the muscle undergoes a bout of chronic regeneration that continues through adulthood *(3, 37)*. By performing Hematoxylin and Eosin (H&E) staining, we confirmed significant myonecrosis in tibialis anterior (TA) muscle of 4-week-old mdx mice and appearance of centronucleated myofibers in 6- and 8-week-old mdx mice (**Fig. 1D**). To investigate the temporal expression of TAK1, we measured levels of TAK1 in skeletal muscle of mdx mice of different age groups. Results showed that levels of TAK1 were significantly increased in GA muscle of 4-week old mdx mice compared to 2-, 6- and 8-week-old mdx mice (**Fig. 1E, F**). In response to stimulation by TNFα, TAK1 phosphorylates receptor interacting protein kinase 1 (RIPK1) at Ser321 residue, which leads to cell death through apoptosis or necroptosis *(38)*. Interestingly, necroptosis mediated myofiber death has been reported to promote muscle degeneration in mdx model of DMD *(39)*. Consistent with increased activation of TAK1, we observed a significant increase in p-RIPK1 (S321), but not total RIPK1 protein levels, in GA muscle of 4-week-old mdx mice compared to other age groups (**Fig. 1E, F)**. To determine the spatial location of TAK1 during the necrotic phase of mdx pathology, we immunostained transverse muscle sections generated from TA muscle of 4-week old mdx mice for TAK1 and Pax7 protein. Results showed that TAK1 was expressed in regenerating myofibers and satellite cells (**Fig. 1G**). Collectively, these results suggest that TAK1 is activated in skeletal muscle of mdx mice during peak necrotic phase.

**Fig. 1.**
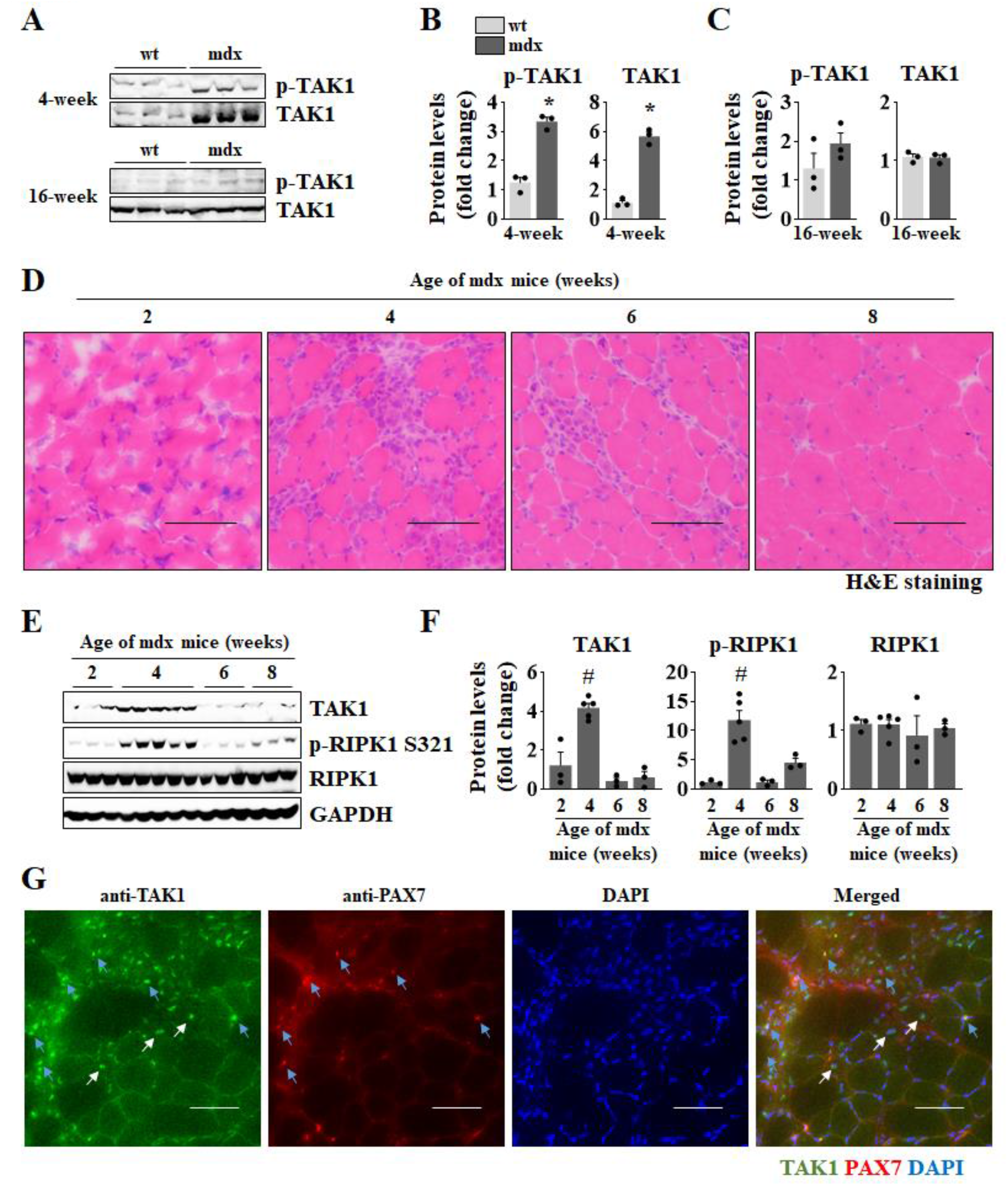
Activation of TAK1 in skeletal muscle of mdx mice. (**A**) Western blot showing the protein levels of p-TAK1 and total TAK1 in GA muscle of 4- and 16-week old wild type (wt) and mdx mice. Quantification of protein levels of p-TAK1 and total TAK1 in **(B**) 4- and **(C)** 16- week old wt and mdx mice. n=3 mice per group. Data represented as mean ± SEM. *p≤0.05, values significantly different from GA muscle of corresponding age-matched wt mice by unpaired Student *t* test. (**D**) Transverse sections of TA muscle from 2-, 4-, 6-, and 8-week-old mdx mice stained with H&E. Scale bar, 100 μm. (**E**) Western blot and (**F**) densitometry analysis showing protein levels of TAK1, p-RIPK1 (S321), total RIPK1 in GA muscle of mdx mice at indicated age. n=3-5. #p≤0.05, values significantly different from GA muscle of 2-week-old mdx mice by one-way ANOVA followed by Tukey’s multiple comparison test. (**G**) Representative images of TA muscle sections from 4-week old mdx mice after immunostaining for TAK1 and Pax7 protein. Scale bar, 50 μm. Blue arrows point to Pax7-positive satellite cells. White arrows point to myonuclei.

### Inactivation of TAK1 causes loss of skeletal muscle mass and contractility in mdx mice

To investigate muscle role of TAK1 in dystrophic mice, we crossed floxed Tak1 (Tak1^fl/fl^) mice with a skeletal muscle-specific tamoxifen inducible Cre line (HSA-MCM) to generate Tak1^mKO^ mice as described *(33, 35)*. The Tak1^mKO^ mice were then crossed with mdx mice to generate mdx;Tak1^fl/fl^ and mdx;Tak1^mKO^ mice. For TAK1 inactivation in skeletal muscle, 3.5-week-old littermate mdx;Tak1^fl/fl^ and mdx;Tak1^mKO^ mice were given daily i.p. injections of tamoxifen (75 mg/kg bodyweight) for 4 consecutive days. The mice were fed a tamoxifen containing chow for the entire duration of the experiments. The mice were analyzed and euthanized after 1 week or 4 weeks of TAK1 inactivation i.e. either at the age of 5 or 8 weeks (**Fig. 2A**). Results showed that TAK1 inactivation in skeletal muscle of mdx mice led to retardation in growth (**Fig. 2B**). The body weight of mdx;Tak1^mKO^ mice was significantly less compared to littermate mdx;Tak1^fl/fl^ mice at the age of 7 and 8 weeks (**Fig. 2B)**. Moreover, 8-week-old mdx;Tak1^mKO^ mice appeared lean and feeble besides having significantly reduced weight gain compared to littermate mdx;Tak1^fl/fl^ mice (**Fig. 2C, D)**. In addition, four paw grip strength of mdx;Tak1^mKO^ mice was significantly reduced compared to mdx;Tak1^fl/fl^ mice (**Fig. 2E**). Next, we investigated muscle contractile properties *in vivo* by measuring the average twitch force, force-frequency relationship, and time to fatigue. Stimulating plantarflexion at different frequencies to estimate the isometric force production showed a significant decrease in average twitch force normalized to body weight in mdx;Tak1^mKO^ mice compared to mdx;Tak1^fl/fl^ mice (**Fig. 2F**). Analysis of force frequency curve showed significantly less force production at frequencies of 50 to 300 Hz in mdx;Tak1^mKO^ mice compared to mdx;Tak1^fl/fl^ mice (**Fig. 2G**). However, there was no significant difference in the fatigability of the muscles between mdx;Tak1^fl/fl^ and mdx;Tak1^mKO^ mice (**Fig. 2H**). We also found reduced GA and TA muscle wet weight in 5- and 8-week-old mdx;Tak1^mKO^ mice compared to corresponding controls (**Fig. 2I, J**). Serum levels of creatine kinase (CK) activity is an important marker of muscle damage in dystrophic mice *(40)*. Results showed a significant decrease in CK levels in 5-week-old mdx;Tak1^mKO^ mice but not 8-week old mdx;Tak1^mKO^ mice compared to their corresponding controls (**Fig. 2K**). These results suggest that muscle-specific inactivation of TAK1 adversely affects muscle mass and contractile properties in mdx mice.

**Fig. 2.**
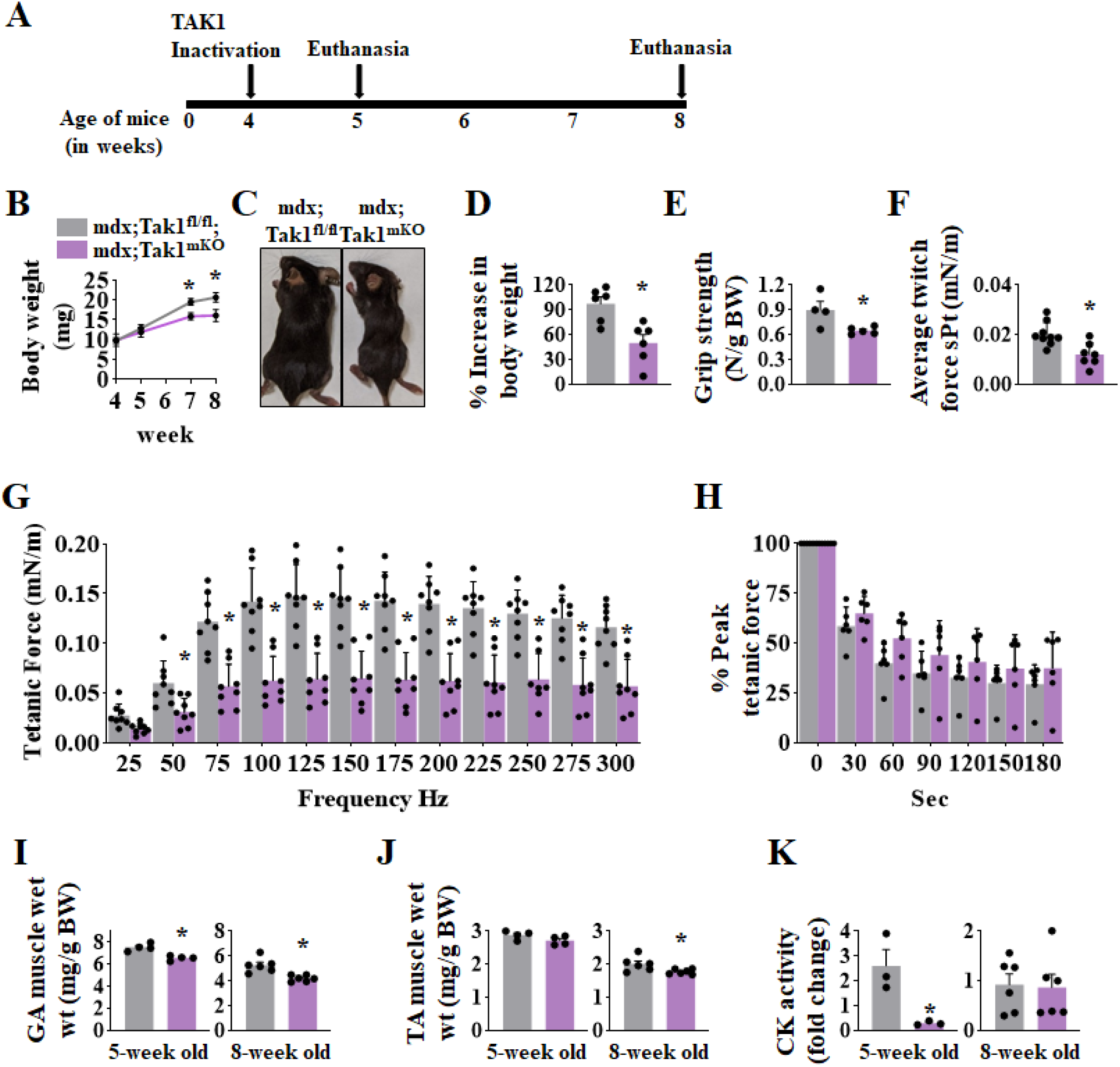
Ablation of TAK1 in young mdx mice blunts skeletal muscle growth and reduce muscle force production. (**A**) Schematic representation showing age of mdx mice, time of TAK1 inactivation and euthanasia for morphometric analysis. (**B**) Average body weight of littermate mdx;Tak1^fl/fl^ and mdx;Tak1^mKO^ mice at indicated ages. (**C**) Gross appearance of mdx;Tak1^fl/fl^ and mdx;Tak1^mKO^ mice at the age of 8 weeks. (**D**) Percentage increase in body weight mdx;Tak1^fl/fl^ and mdx;Tak1^mKO^ mice with age. (**E**) Average four paw grip strength per gram of body weight of 8-week-old mdx;Tak1^fl/fl^ and mdx;Tak1^mKO^ mice. n=4-6. Quantification of normalized (**F**) average specific twitch force, (**G**) tetanic forces to stimulation frequency relationship, and (**H**) peak tetanic force over 180 sec in 8-week-old mdx;Tak1^fl/fl^ and mdx;Tak1^mKO^ mice. n=7-9. Wet weight of (**I**) GA and (**J**) TA muscle and (**K**) Fold change in serum levels of CK activity in mdx;Tak1^fl/fl^ and mdx;Tak1^mKO^ mice at 5- and 8-week of age. n=3-6. Data represented as mean ± SEM. *p≤0.05, values significantly different from corresponding mdx;Tak1^fl/fl^ mice by unpaired Student *t* test by one-way ANOVA followed by Tukey’s multiple comparison test.

### Targeted inactivation of TAK1 in young mdx mice reduces muscle injury but blunts myofiber growth

To understand the effects of inactivation of TAK1 on muscle histopathology, we generated transverse sections of diaphragm, TA, and soleus muscle isolated from 5-or 8-week-old mdx;Tak1^fl/fl^ and mdx;Tak1^mKO^ mice. H&E staining of muscle sections showed considerable necrotic area and cellular infiltration in skeletal muscle of 5-week-old mdx;Tak1^fl/fl^ mice (**Fig. 3A**). At the age of 8 weeks, the necrotic areas in diaphragm, TA, and soleus muscle sections were filled with regenerating myofibers with central nucleation in mdx;Tak1^fl/fl^ mice (**Fig. 3B)**. Intriguingly, these histopathological features were considerable reduced in skeletal muscle of mdx;Tak1^mKO^ mice (**Fig. 3A, B**). Quantitative analysis showed that the proportion of centronucleated myofibers (CNF) in the TA and soleus muscle of 5-week-old mdx;Tak1^mKO^ mice was significantly reduced compared to age matched mdx;Tak1^fl/fl^ mice (**Fig. 3C**). Moreover, diaphragm, TA, and soleus muscle of 8-week-old mdx;Tak1^mKO^ mice also showed significantly reduction in number of CNFs compared to littermate mdx;Tak1^fl/fl^ mice (**Fig. 3D**). Interestingly, the myofibers of mdx;Tak1^mKO^ mice appeared to be considerably smaller in size compared to mdx;Tak1^fl/fl^ mice (**Fig. 3A, B**). Indeed, quantitative analysis showed a significant reduction in average myofiber cross-sectional area (CSA) in diaphragm and TA muscle of 5-week-old mdx;Tak1^mKO^ mice compared to littermate mdx;Tak1^fl/fl^ mice (**Fig. 3E**). Correspondingly, average myofiber CSA in diaphragm, TA and soleus muscle was also significantly reduced in 8-week-old mdx;Tak1^mKO^ mice compared to littermate controls (**Fig. 3F**). This phenotype was more prominent at the age of 8 weeks, where average myofiber CSA in TA and soleus muscle was reduced by more than 50% with TAK1 inactivation.

**Fig. 3.**
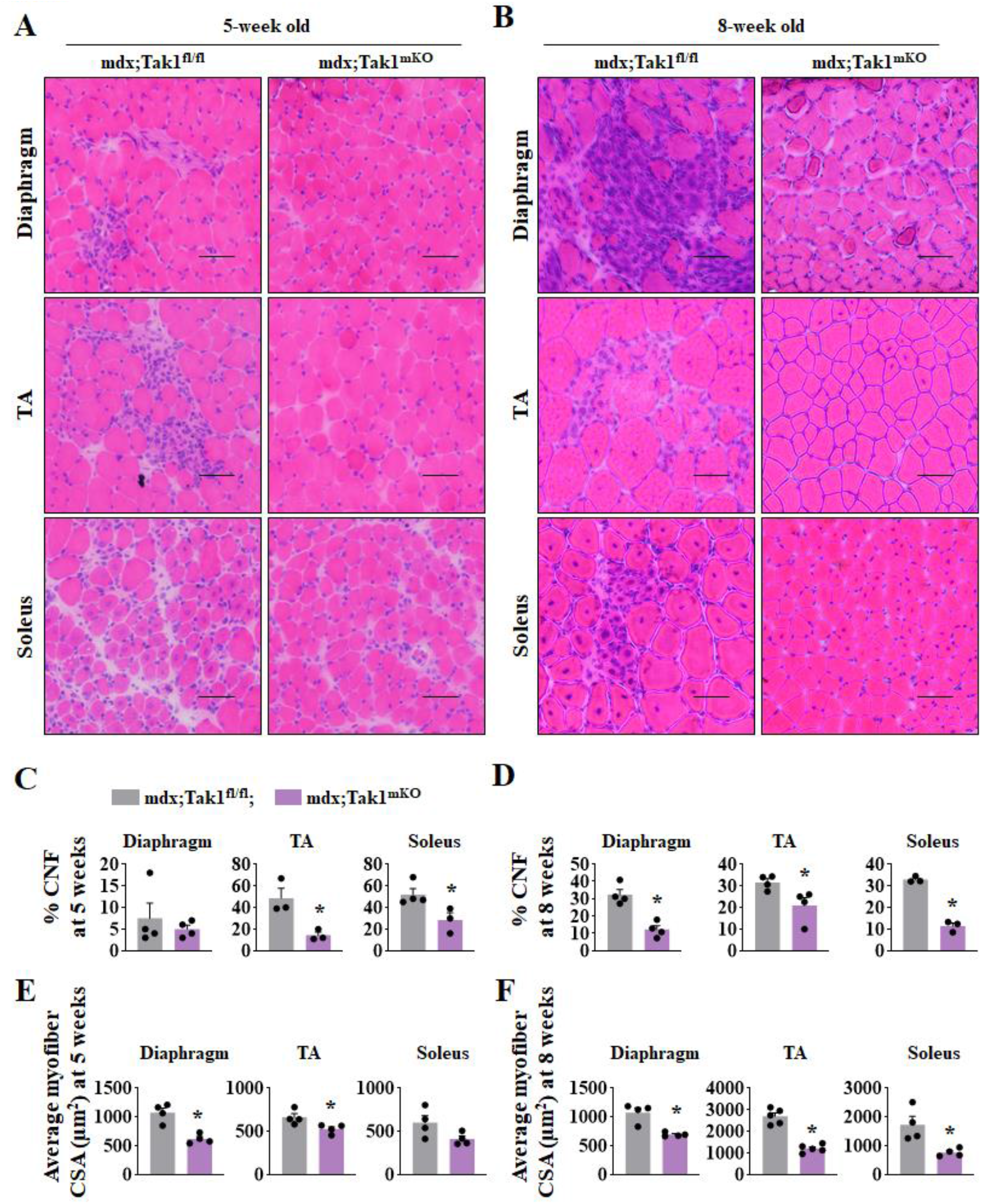
TAK1 deletion improves muscle histopathology but suppresses myofiber growth in young mdx mice. 3.5 week old mdx;Tak1^fl/fl^ and mdx;Tak1^mKO^ mice were treated with tamoxifen and analyzed at the age of 5 or 8 weeks. Representative photomicrographs of transverse sections of diaphragm, TA and soleus muscle of (**A**) 5-week (**B**) 8-week old mdx;Tak1^fl/fl^ and mdx;Tak1^mKO^ mice after staining with H&E. Scale bar, 50 μm. Quantification of percentage of CNF in diaphragm, TA and Soleus muscle of (**C**) 5-week and (**D**) 8-week old mdx;Tak1^fl/fl^ and mdx;Tak1^mKO^ mice. Average myofiber cross-section area (CSA) of diaphragm, TA and Soleus muscle of (**E**) 5-week and (**F**) 8-week old mdx;Tak1^fl/fl^ and mdx;Tak1^mKO^ mice. n=3-5. Data represented as mean ± SEM. *p≤0.05, values significantly different from corresponding mdx;Tak1^fl/fl^ mice by unpaired Student *t* test

### TAK1 contributes to myofiber damage through necroptosis in mdx mice

Mutations in the dystrophin gene destabilizes the dystrophin-glycoprotein complex and causes severe myofiber necrosis in skeletal muscle *(41)*. A recent study has shown that programmed necrosis or necroptosis is pronouncedly activated in the mdx model of DMD and is responsible for RIPK1-dependent myofiber death *(39)*. Sustained phosphorylation at the intermediate domain of RIPK1 (S321) by TAK1 has been demonstrated to trigger necroptosis *(38)*. Therefore, we sought to investigate whether TAK1 deletion in the pre-necrotic skeletal muscle of mdx mice can suppress myonecrosis by inhibiting RIPK1 phosphorylation. Indeed, western blot analysis showed a significant reduction in p-RIPK1 (S321) levels in 5- and 8-week-old mdx;Tak1^mKO^ mice compared to corresponding controls (**Fig. 4A, B**). To investigate the extent of necrosis we transverse sections of diaphragm, TA and soleus muscle from 8-week-old mdx;Tak1^fl/fl^ and mdx;Tak1^mKO^ mice were stained with anti-mouse IgG antibody. Significantly larger IgG-filled necrotic areas were observed in mdx;Tak1^fl/fl^ mice compared to mdx;Tak1^mKO^ mice (**Fig. 4C, D**). These results suggest that inactivation of TAK1 limits muscle damage by inhibiting RIPK1 phosphorylation.

**Fig. 4.**
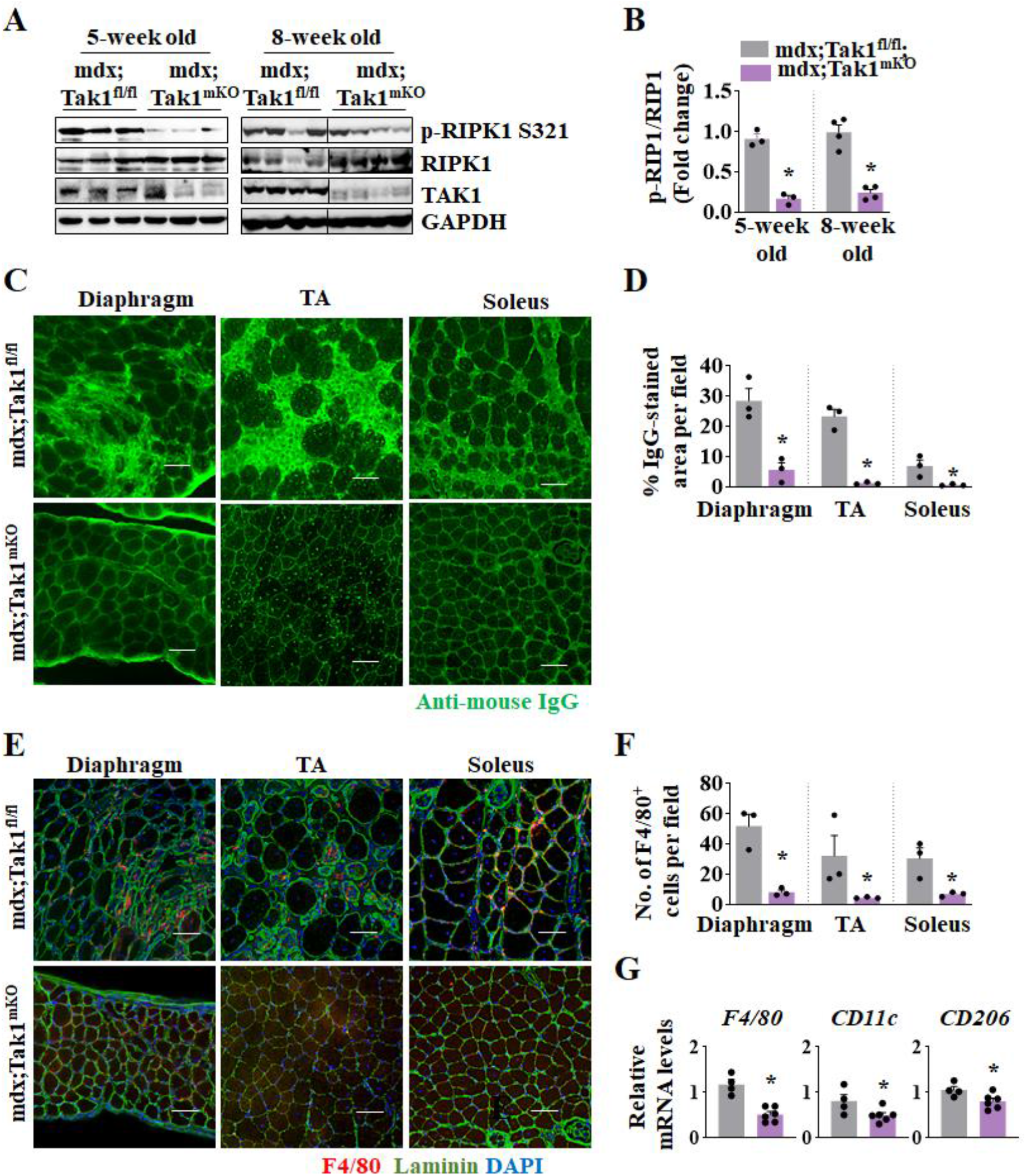
TAK1 contributes to myonecrosis in dystrophic muscle through RIPK1 phosphorylation. (**A**) Western blot showing protein levels of p-RIPK1 (S321), total RIPK1, and TAK1 in GA muscle of 5- and 8-week old mdx;Tak1^fl/fl^ and mdx;Tak1^mKO^ mice. **(B)** Densitometry analysis showing the ratio of p-RIPK1 to total RIP in GA muscle of 5-week and 8- week old mdx;Tak1^fl/fl^ and mdx;Tak1^mKO^ mice. n=3-4. (**C**) Representative photomicrographs of transverse sections of diaphragm, TA and soleus muscle of 8-week-old mdx;Tak1^fl/fl^ and mdx;Tak1^mKO^ mice stained with anti-mouse IgG antibody. Scale bar, 50 μm. (**D**) Percentage of IgG-stained area per field in diaphragm, TA and soleus muscle of 8-week-old mdx;Tak1^fl/fl^ and mdx;Tak1^mKO^ mice. (**E**) Representative photomicrographs of transverse sections of diaphragm, TA, and soleus muscle of 8-week-old mdx;Tak1^fl/fl^ and mdx;Tak1^mKO^ mice immunostained for F4/80 antigen and Laminin protein. DAPI was used to visualize nuclei. **(F)** Quantification of F4/80-postive cells per field in diaphragm, TA, and soleus muscle of 8-week-old mdx;Tak1^fl/fl^ and mdx;Tak1^mKO^ mice. n=3. (**G**) Relative mRNA levels of F4/80, CD11c, and CD206 in GA muscle of 8-week old mdx;Tak1^fl/fl^ and mdx;Tak1^mKO^ mice. n=4-6 mice per group. Data represented as mean ± SEM. *p≤0.05, values significantly different from mdx;Tak1^fl/fl^ mice by unpaired Student *t* test.

In response to myofiber injury, inflammatory immune cells, such as macrophages infiltrate into dystrophic muscle *(42, 43)*. Immunostaining for F4/80 antigen, a marker for murine macrophages, showed a significant decrease in the number of F4/80^+^ cells in diaphragm, TA, and soleus muscle of 8-week-old mdx;Tak1^mKO^ mice compared to littermate mdx;Tak1^fl/fl^ mice (**Fig. 4E,F**). Inflammatory response elicits a sequential activation and homing of M1 and M2 macrophages, which play a critical role in skeletal muscle degeneration and regeneration *(44, 45)*. CD11c^+^ proinflammatory M1 macrophages infiltrate the dystrophic muscle and contributes to myolysis whereas during the regenerative phase, a shift towards CD206-expressing anti-inflammatory M2 macrophages facilitates repair *(45-47)*. We found a significant reduction in the relative mRNA levels of F4/80, CD11c, and CD206 in the GA muscle of 8-week-old mdx;Tak1^mKO^ mice compared to littermate mdx;Tak1^fl/fl^ mice (**Fig. 4G**). These results suggest an active inflammatory and regenerative phase in mdx;Tak1^fl/fl^ mice that has subsided in mdx;Tak1^mKO^ mice.

### TAK1 regulates autophagy and muscle regeneration in mdx mice

Repressed autophagy and mitophagy response are a major manifestation of DMD pathology *(10, 48, 49)*. We investigated whether TAK1 regulates autophagy and mitophagy in dystrophic muscle of mdx mice. Our analysis showed a significant increase in the relative mRNA levels of mitophagy regulator Pink1, but not Park2, in both 5- and 8-week-old mdx;Tak1^mKO^ mice compared to corresponding controls (**Fig. 5A**). Bnip3 is an important regulator of autophagy as well as mitophagy *(50)*. We found a significant increase in mRNA levels of Bnip3 in 5-week-old mdx;Tak1^mKO^ mice compared to littermate controls (**Fig. 5B**). In addition, the mRNA levels of LC3B, Atg12, and Gabarpl1, but not Atg5 and Beclin1, were significantly up-regulated in GA muscle of mdx;Tak1^mKO^ mice compared to littermate mdx;Tak1^fl/fl^ mice (**Fig. 5C**). Finally, western blot and densitometry analysis showed a significant increase in the ratio of LC3BII/LC3BI, a classical marker of autophagy, in dystrophic muscle of 8-week-old mdx;Tak1^mKO^ mice compared to mdx;Tak1^fl/fl^ mice (**Fig. 5D, E**). These results suggest that inactivation of TAK1 leads to the activation of autophagy and mitophagy in dystrophic muscle of mdx mice.

**Fig. 5.**
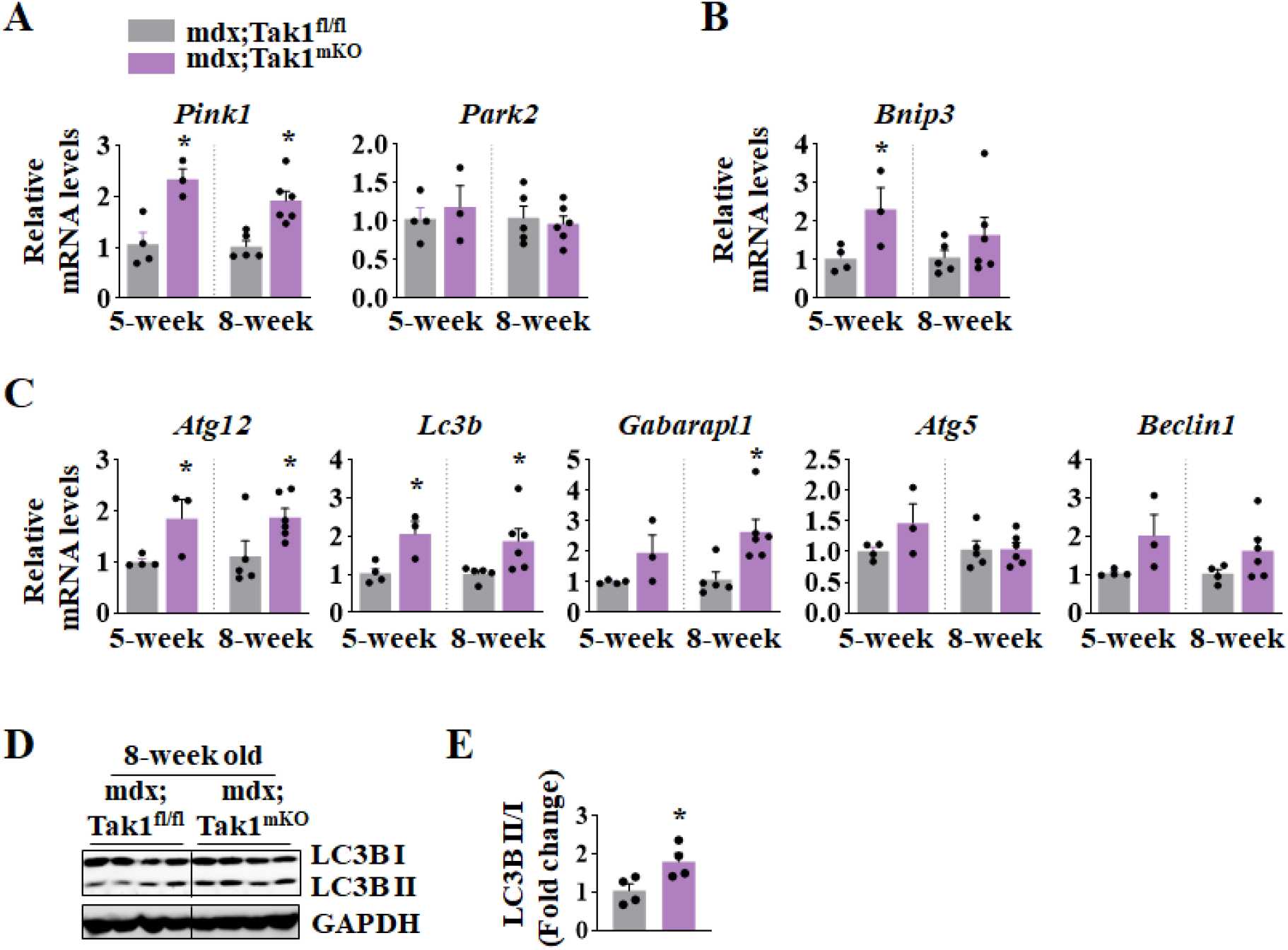
Inactivation of TAK1 improves markers of mitophagy and autophagy in dystrophic muscle of mdx mice. 3.5-week old mdx;Tak1^fl/fl^ and mdx;Tak1^mKO^ mice were treated with tamoxifen and analyzed at the age of 5 or 8 weeks by qRT-PCR and western blotting. Relative mRNA levels of mitophagy markers (**A**) *Pink1* and *Park2*, (**B**) *Bnip3* and autophagy markers (**C**) *Atg12, LC3b*, Gabarapl1, *Atg5* and *Beclin1* in GA muscle of 5-or 8-week-old mdx;Tak1^fl/fl^ and mdx;Tak1^mKO^ mice. n=3-6. (**D**) Western blot and (**E**) densitometry analysis showing the ratio of LC3BII to LC3BI in GA muscle of 8-week old mdx;Tak1^fl/fl^ and mdx;Tak1^mKO^ mice. n=4-6. Data represented as mean ± SEM. *p≤0.05, values significantly different from mdx;Tak1^fl/fl^ mice by unpaired Student *t* test.

Exhaustion of satellite stem cells and blunted myogenic potential with increasing age is a manifestation of DMD *(51)*. Intriguingly, a few recent studies have suggested that depleting myogenic progenitor cells can ameliorate myopathy severity in animal models of muscular dystrophies *(52, 53)*. Therefore, we sought to investigate whether TAK1 deletion in dystrophic muscle affects regenerative myogenesis process. For this analysis, 3.5-week old mdx;Tak1^fl/fl^ and mdx;Tak1^mKO^ mice were treated with tamoxifen and the mice were analyzed at the age of 8 weeks. Results showed no significant difference in mRNA or protein levels of Pax7 in GA muscle of mdx;Tak1^fl/fl^ and mdx;Tak1^mKO^ mice (**Supplementary Fig. S1A-C**). Immunostaining for the embryonic isoform of myosin heavy chain (eMyHC) showed that the newly formed eMyHC expressing myofibers in GA muscle sections were significantly smaller in size and fewer in number in mdx;Tak1^mKO^ mice compared to GA mdx;Tak1^fl/fl^ mice (**Supplementary Fig. S1D, E, F**). Next, we measured the mRNA levels of some myogenic regulatory factors. Results showed no significant difference in the relative mRNA levels of eMyHC (Myh3) and Myf5 in GA muscles of mdx;Tak1^mKO^ and mdx;Tak1^fl/fl^ mice (**Supplementary Fig. S1G**). However, we found a significant increase in mRNA levels of MyoD and Myogenin in GA muscle of mdx;Tak1^mKO^ mice compared to mdx;Tak1^fl/fl^ mice (**Supplementary Fig. S1G**).

Notch and Wnt signaling pathways play critical roles in embryonic development of skeletal muscle and regenerative myogenesis *(54-56)*. To gain further insight into the role of TAK1 in the regulation of Notch and Wnt pathways in dystrophic muscle of mdx mice we analyzed the mRNA levels of a few key components of these pathways. Results showed a significant increase in the mRNA levels of Notch ligands *Jagged1* and *Jagged 2*, Notch receptors *Notch2* and *Notch3*, and Notch targets *Hes1, Hes6, Hey1*, and *HeyL* in GA muscle of 8-week old mdx;Tak1^mKO^ mice compared to littermate mdx;Tak1^fl/fl^ mice. In contrast, there was no significant difference in mRNA levels of *Dll1, Dll4*, and *Notch1* in GA muscle of mdx;Tak1^mKO^ mice compared to mdx;Tak1^fl/fl^ mice (**Supplementary Fig. S2A, B, C**). Next, we measured mRNA levels of Wnt ligands (Wnt3a, Wnt4, Wnt5a, Wnt7a, and Wnt11), receptors (Fzd1, Fzd2, Fzd4 and Fzd6) and target gene (Axin2) of Wnt signaling pathway. There was a significant increase in mRNA levels of *Wnt5a, Wnt11, Fzd4*, and *Axin2* in GA muscle of mdx;Tak1^mKO^ mice compared to littermate mdx;Tak1^fl/fl^ mice (**Supplementary Fig. S2D, E, F**). These results suggest that deletion of TAK1 leads to the activation Notch and Wnt signaling in dystrophic muscle of mdx mice.

### Targeted inactivation of TAK1 in post-necrotic mdx mice causes muscle wasting

We have previously reported that TAK1 is essential for maintaining skeletal muscle mass in adult wild-type mice *(33)*. However, the role of TAK1 in the regulation of skeletal muscle mass at the post-necrotic phase in mdx mice remains unknown. Therefore, we inactivated TAK1 in 12-week-old adult littermate mdx;Tak1^fl/fl^ and mdx;Tak1^mKO^ mice and performed morphometric and biochemical analysis at the age of 16 weeks (**Fig. 6A**). Results showed a significant reduction in bodyweight of mdx;Tak1^mKO^ mice compared to littermate mdx;Tak1^fl/fl^ mice (**Fig. 6B**). Next, we investigated whether TAK1 deletion affects force production in skeletal muscle of adult mdx mice. A significant reduction in specific twitch force normalized to body weight was observed in mdx;Tak1^mKO^ mice compared to mdx;Tak1^fl/fl^ mice (**Fig. 6C**). Moreover, there was significant reduction in tetanic force production with stimulation ranging from 50-300 Hz in mdx;Tak1^Mko^ mice compared to littermate controls (**Fig. 6D**). Finally, the mice were euthanized and hind limb muscles were isolated for morphometric analysis. We found a significant reduction in wet weight of TA and GA muscles compared to corresponding mdx;Tak1^fl/fl^ mice (**Fig. 6E, F**). However, serum levels of CK remained comparable between mdx;Tak1^fl/fl^ and mdx;Tak1^mKO^ mice (**Fig. 6G**). By performing western blot, we confirmed a drastic reduction in the levels of TAK1 protein in skeletal muscle of mdx;Tak1^mKO^ mice compared littermate mdx;Tak1^fl/fl^ mice (**Fig. 7H**).

**Fig. 6.**
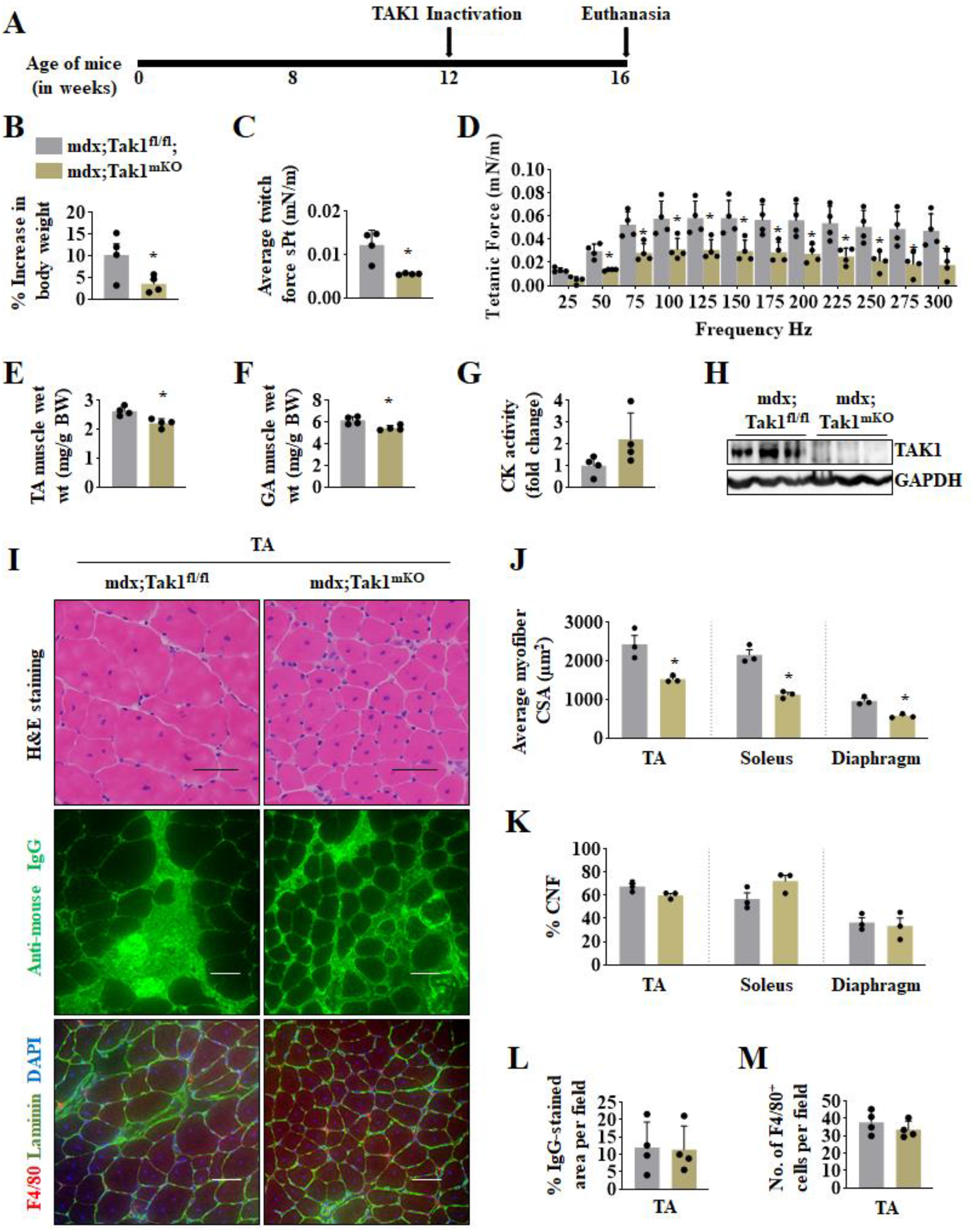
Targeted ablation of TAK1 in adult mdx mice causes muscle wasting without affecting histopathology. (**A**) Schematic representation showing the age of mice and time of treatment with tamoxifen and euthanasia. (**B**) Percentage change in body weight from initial weight of mdx;Tak1^fl/fl^ and mdx;Tak1^mKO^ mice. (**C**) Normalized average specific twitch force and (**D**) tetanic force with increasing stimulation frequency in 16-week old mdx;Tak1^fl/fl^ and mdx;Tak1^mKO^ mice. Wet weight of (**E**) TA, and (**F**) GA muscle normalized with body weight **(G)** serum levels of CK in 16-week old mdx;Tak1^fl/fl^ and mdx;Tak1^mKO^ mice. n=4. (**H**) Western blot showing protein levels of TAK1 in GA muscle of mdx;Tak1^fl/fl^ and mdx;Tak1^mKO^ mice. (**I**) Representative photomicrographs of TA muscle sections stained with H&E, anti-mouse IgG, or co-immunostained for F4/80, Laminin and DAPI. (**J**) Average myofiber cross-section area and (**K**) Percentage of CNF in diaphragm, TA, and soleus muscle of mdx;Tak1^fl/fl^ and mdx;Tak1^mKO^ mice. n=3. Quantitative analysis of (**L**) percentage of IgG-stained area per field, and (**M**) F4/80-positive cells per field in TA muscle of 16-week-old mdx;Tak1^fl/fl^ and mdx;Tak1^mKO^ mice. n=3. Data represented as mean ± SEM. *p≤0.05, values significantly different from GA muscle of 16-week-old mdx;Tak1^fl/fl^ mice by unpaired Student *t* test.

**Fig. 7.**
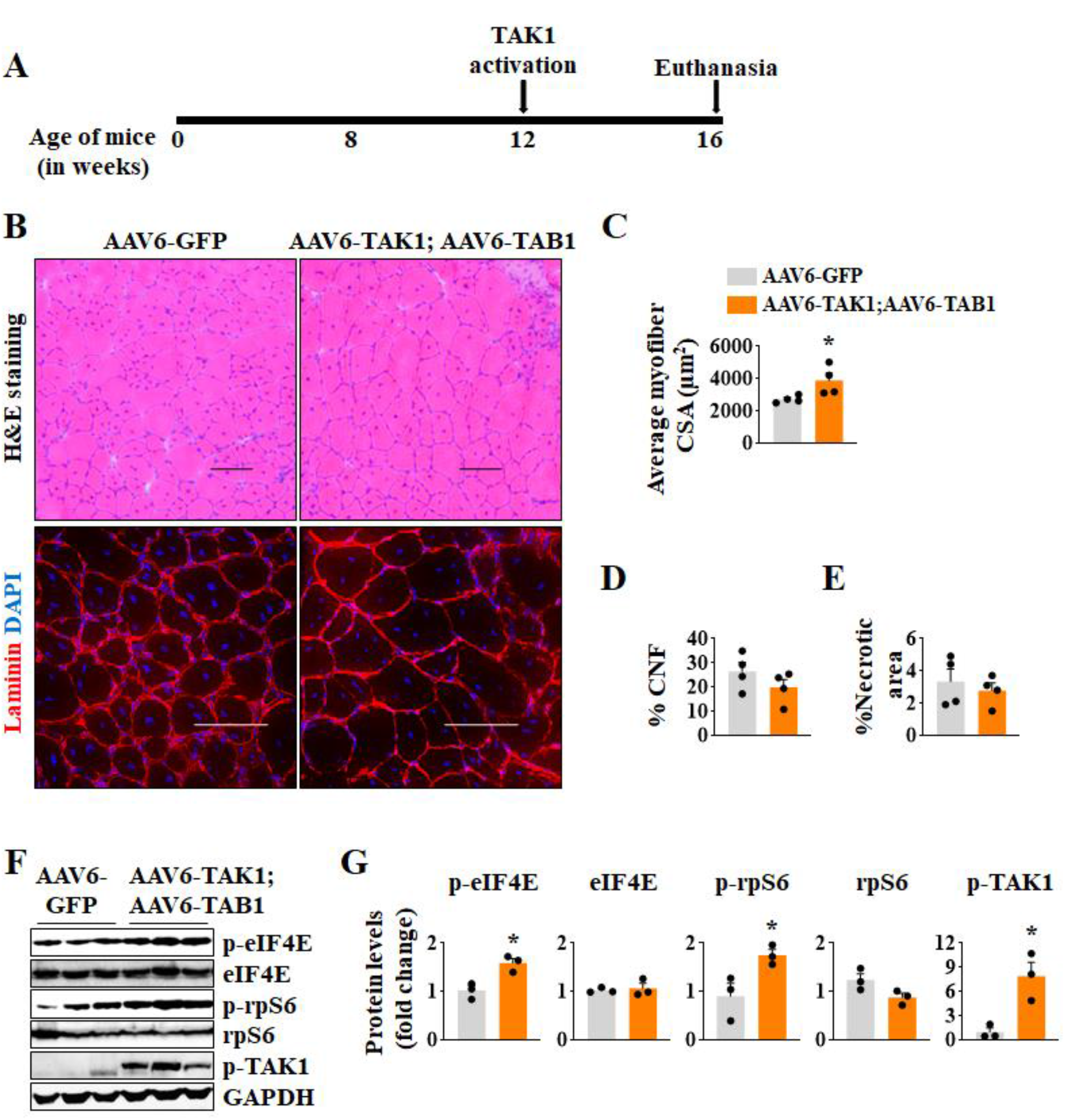
TAK1 activation in adult mdx mice promotes myofiber growth. (**A**) Schematic representation showing the age of mdx mice, time of AAVs injection, and euthanasia. (**B**) Representative photomicrographs showing transverse sections after stained with H&E or immunostaining with Laminin and DAPI. Scale bar, 100μm. Quantitative analysis for (**C**) Average myofiber cross-section area, (**D**) Percentage of centronucleated fibers, and (**E**) Percentage of necrotic area in TA muscle of mdx mice injected with AAV6-GFP or a combination of AAV6-TAK1 and AAV6-TAB1. n=4. (**F**) Western blot and (**G**) densitometry analysis showing protein levels of p-eIF4E, eIF4E, p-rpS6, rpS6, p-TAK1, and unrelated protein GAPDH in TA muscle of mdx mice injected with AAV6-GFP or co-injected with AAV6-TAK1 and AAV6-TAB1. n=3. Data represented as mean ± SEM. *p≤0.05, values significantly different from TA muscle of mdx mice injected with AAV6-GFP.

To investigate the effect of inactivation of TAK1 on muscle histopathology in post-necrotic mdx mice, transverse sections of diaphragm, TA, and soleus muscle were processed for H&E staining. Results showed a significant reduction in average myofiber CSA in skeletal muscle of mdx;Tak1^mKO^ mice compared to corresponding controls (**Fig. 6I, J; Supplementary Fig. S3A**). However, the proportion of centronucleated myofibers remained comparable between mdx;Tak1^fl/fl^ and mdx;Tak1^mKO^ mice (**Fig. 6I, K; Supplementary Fig. S3A**). Additionally, we did not find any difference in area under necrosis between mdx;Tak1^mKO^ and mdx;Tak1^fl/fl^ mice analyzed by staining of muscle sections with anti-mouse IgG (**Fig. 6I, L; Fig. S3B**). To investigate whether muscle-specific TAK1 inactivation affects macrophage infiltration in dystrophic muscle of adult mdx mice, we immunostained muscle sections for F4/80 antigen. Results showed that number of F4/80^+^ cells was comparable between mdx;Tak1^fl/fl^ and mdx;Tak1^mKO^ mice (**Fig.6I, M; Supplementary Fig. S3C**).

We also performed Sirius red staining to measure collagen levels in dystrophic muscle. However, there was no apparent difference in collagen deposition in diaphragm of 16-week old mdx;Tak1^fl/fl^ and mdx;Tak1^mKO^ mice (**Supplementary Fig. S3D**). Moreover, mRNA levels of Col1a1, Col2a1, Col3a1 and Col4a1 remained comparable in GA muscle of mdx;Tak1^fl/fl^ and mdx;Tak1^mKO^ mice (**Supplementary Fig. S3E**). Collectively, these results suggest that inactivation of TAK1 in adult mdx mice leads to loss of skeletal muscle mass without affecting tissue histopathology.

### Forced activation of TAK1 promotes myofiber growth in adult mdx mice

We next sought to determine the effect of forced activation of TAK1 in skeletal muscle of adult mdx mice. TAK1 forms a complex with adaptor protein TAB1, which is essential for TAK1 autophosphorylation at S192 and subsequent activation *(30, 57)*. We co-expressed TAK1 and TAB1 in TA muscle of 12-week old mdx mice using AAV serotype 6 (AAV6) vectors having a constitutive cytomegalovirus (CMV) promoter. AAV6-GFP vector was used as a control similar to as described *(35)*. Our initial analysis showed that intramuscular injection of about 2 × 10^10^ vgs of AAV6-GFP vectors in TA muscle of mdx mice results in GFP expression in almost all myofibers at day 28 post injection (**Supplementary Fig. S4**). Therefore, we transduced TA muscle of 12-week old mdx mice by injecting AAV6-TAK1 (1.25 × 10^10^ vg) and TAB1 (1.25 × 10^10^ vg). The contralateral TA muscle was injected with AAV6-GFP (2.5 × 10^10^ vg) vector. After 28 days, the mice were euthanized and the TA muscle was analyzed by morphometric and biochemical assays (**Fig. 7A**). H&E staining of muscle sections did not show any overt histological changes in TA muscle co-injected with AAV6-TAK1 and AAV6-TAB1 or with AAV6-GFP alone. The necrotic areas remained filled with centronucleated myofibers, a typical feature in skeletal muscle of adult mdx mice (**Fig. 7B**). Anti-laminin staining followed by quantitative analysis showed that average myofiber CSA was significantly higher in TA muscle injected with AAV6-TAK1 and AAV6-TAB1 compared to contralateral TA muscle injected with AAV6-GFP alone (**Fig. 7B, C**). However, empty necrotic areas filled with cell infiltrate and the proportion of centronucleated myofibers remained comparable between TA muscle injected with AAV6-GFP alone or with a combination of AAV6-TAK1 and AAV6-TAB1 (**Fig. 7B, D, E**). To understand the mechanism of myofiber hypertrophy in mdx mice with TAK1 activation, we measured the phosphorylation levels of proteins that positively regulate skeletal muscle mass. We found a significant increase in the levels of phosphorylated rpS6 (p-rpS6), phosphorylated eIF4E (p-eIF4E) and phosphorylated TAK1 (p-TAK1) in TA muscle overexpressing TAK1 and TAB1 compared to GFP alone (**Fig. 7F, G**) suggesting that forced activation TAK1 causes myofiber growth potentially through stimulating protein synthesis in mdx mice.

### Temporal regulation of TAK1 expression improves muscle histopathology

Considering our preceding results, we speculated that expression of *Tak1* gene is temporally regulated with age in dystrophic muscle of mdx mice. Upregulated *Tak1* expression exacerbates injury during the initial myonecrotic phase and once the necrosis-regenerative process stabilizes to lower rate around 6-8 weeks, TAK1 functions to maintain and support muscle growth. To validate our hypothesis, we devised a strategy to inactivate TAK1 at the onset of necrosis-regeneration phase and restore TAK1 activity during the post necrotic regenerative phase. For this experiment, we used littermate mdx;Tak1^fl/fl^ and mdx;Tak1^mKO^ mice and inactivated TAK1 at the age of 4 weeks by tamoxifen treatment. After two weeks of TAK1 deletion, TA muscle of littermate mdx;Tak1^fl/fl^ and mdx;Tak1^mKO^ mice was injected with AAV6-TAK1 (2.5 × 10^10^ vg) and AAV6-TAB1 (2.5 × 10^10^ vg). Contralateral TA muscle was injected with control AAV6-GFP vector (2.5 × 10^10^ vg). After 4 weeks (i.e. at the age of 10 weeks), the mice were euthanized and the TA muscle was isolated for morphometric analysis (**Fig. 8A**). H&E staining showed that TA muscle from mdx;Tak1^mKO^ mice had substantially fewer centronucleated fibers and less empty areas filled with cellular infiltrates compared to TA muscle from mdx;Tak1^fl/fl^ mice (**Fig. 8B**). Co-injection of AAV6-TAK1 and AAV6-TAB1 or with AAV6-GFP alone, did not cause any noticeable change in these features in mdx;Tak1^fl/fl^ or mdx;Tak1^mKO^ mice (**Fig. 8B**). To estimate the extent of necrotic areas, we performed immunostaining with Alexa 568-labelled mouse IgG. Results confirmed a significantly lesser numbers of IgG filled myofibers in both GFP-expressing or TAK1- and TAB1-expressing TA muscle of mdx;TAK1^mKO^ mice compared to corresponding GFP-expressing or TAK1- and TAB1-expressing TA muscle of mdx;Tak1^fl/fl^ mice (**Fig. 8B, C**). Additionally, estimating the centronucleation showed a significant reduction in the proportion of centronucleated myofibers in both GFP-expressing or TAK1- and TAB1-expressing TA muscle of mdx;TAK1^mKO^ mice compared to corresponding GFP-expressing or TAK1- and TAB1-expressing TA muscle of mdx;Tak1^fl/fl^ mice (**Fig. 8B, D**). However, a significant increase in average myofiber CSA was noticeable in TA muscle co-injected with AAV6-TAK1 and AAV6-TAB1 compared to TA muscle injected with AAV6-GFP alone in mdx;Tak1^fl/fl^ mice. Moreover, in mdx;Tak1^mKO^ mice, the average myofiber CSA of TAK1- and TAB1-overexpressing TA muscle was significantly higher than GFP-expressing TA muscle (**Fig. 8B, E**). Skeletal muscle of mdx mice contains myofibers with varying size, some with unusually large myofiber diameter than wild-type mice. Therefore, we compared the myofiber size distribution between TAK1- and TAB1-expressing TA muscle from mdx;Tak1^mKO^ mice and GFP-expressing mdx;Tak1^fl/fl^ mice with TA muscle of age-matched wild-type mice (**Supplementary Fig. S5A**). Relative frequency distribution analysis showed that myofiber CSA of TA muscle from mdx;Tak1^mKO^ mice overexpressing TAK1 and TAB1 was similar to that of TA muscle of age matched wild-type mice (**Supplementary Fig. S5B**). Conversely, a proportion of myofibers in the TA muscle of GFP-expressing mdx;Tak1^fl/fl^ mice had abnormally large CSA compared to myofibers from TA muscle of age-matched wild-type mice (**Supplementary Fig. S5B**). Altogether, these results suggest that tuning TAK1 expression with age alleviates dystrophinopathy in mdx mice.

**Fig. 8.**
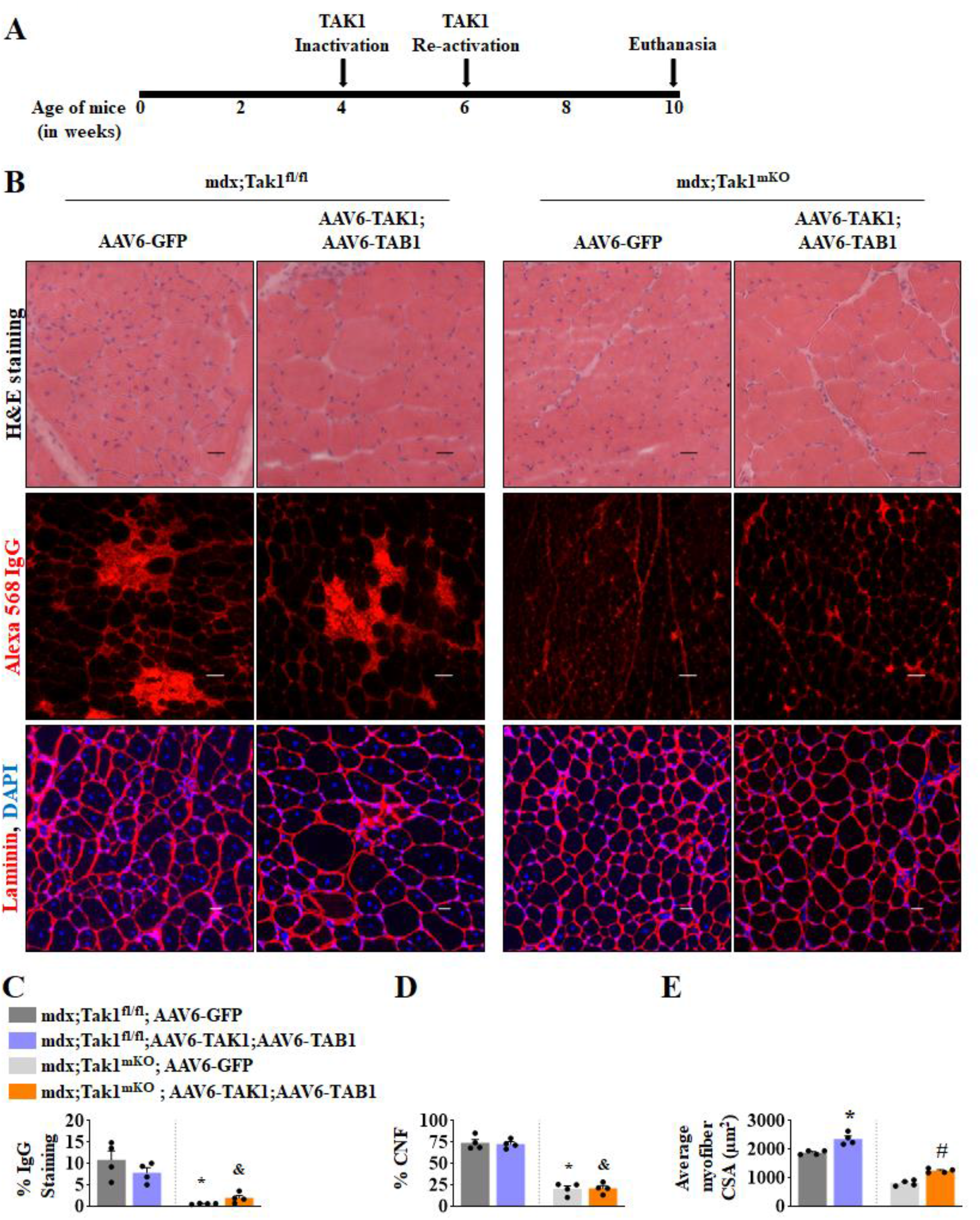
Temporal control of TAK1 expression in dystrophic muscle ameliorates histopathology. (**A**) Schematic representation showing the age of mdx mice, time of TAK1 inactivation, AAV injection, and euthanasia. (**B**) Representative photomicrographs showing TA muscle sections after H&E staining, anti-mouse IgG staining, or co-immunostained for Laminin and DAPI. Scale bar, 50 μm. (**C**) Percentage of IgG-filled necrotic area, (**D**) percentage of centronucleated fibers (CNFs), and (**E**) Average myofiber cross-section area in TA muscle of mdx;Tak1^fl/fl^ and mdx;Tak1^mKO^ mice injected with AAV6-GFP or a combination of AAV6-TAK1 and AAV6-TAB1. n=4. Data represented as mean ± SEM and was analyzed by two-way ANOVA followed by Tukey’s multiple comparison test. *p≤0.05, values significantly different from TA muscle of mdx;Tak1^fl/fl^ injected with AAV6-GFP. ^&^p≤0.05, values significantly different from TA muscle of mdx;Tak1^fl/fl^ mice co-injected with AAV6-TAK1 and AAV6-TAB1. #p≤0.05, values significantly different from TA muscle of mdx;Tak1^mKO^ mice injected with AAV6-GFP.

## DISCUSSION

Mutation in dystrophin gene is present at birth, yet most DMD cases are diagnosed at 2-3 years of age when symptoms emerge *(58)*. However, the molecular and signaling mechanisms that trigger onset and progression of myopathy in DMD patients remain less understood. In this study, we have investigated the role of TAK1 in dystrophinopathy. Our results demonstrate that TAK1 is an important regulator of dystrophic phenotype in mdx mice. TAK1 is highly activated in the skeletal muscle of young mdx mice during the initial necrotic spike where its role is to accelerate myofiber injury and augment inflammation and myofiber necroptosis. Conversely, TAK1 also functions to maintain skeletal muscle mass and is indispensable for skeletal muscle growth in both and young and adult mdx mice. Our results demonstrate that forced activation of TAK1 promotes myofiber growth without producing any adverse effects on muscle histopathology in mdx mice.

TAK1 is an important signaling protein that mediates the activation of multiple downstream signaling pathways, including NF-κB and p38 MAPK in response to receptor-mediated events *(27, 31, 59)*. Previous studies have shown that NF-κB is activated in skeletal muscle of mdx mice and targeted inhibition of NF-κB in myofibers or macrophages improves dystrophic phenotype in mdx mice *(18-20, 60)*. Similarly, it has been reported that the activation of p38MAPK contributes to myofiber injury in dystrophic muscle of mdx mice *(21, 22)*. Consistent with its role in the activation of NF-κB and p38MAPK, we found that targeted inducible inactivation of TAK1 reduces muscle injury and accumulation of macrophages in dystrophic muscle of mdx mice (**Figs. 3, 4**). However, we did not find any improvement in muscle mass or function. Indeed, our experiments demonstrated that skeletal muscle mass and function are significantly reduced upon inactivation of TAK1 in both young and adult mdx mice (**Figs. 2, 6**). These results suggest that while TAK1 contributes to inflammation and myonecrosis during necrotic phase, it is also essential for myofiber growth in dystrophic muscle of mdx mice. Intriguingly, our results also demonstrate that forced activation of TAK1 through AAV-mediated overexpression of TAK1 and TAB1 does not produce any adverse effects on muscle histopathology but promotes growth of myofibers in adult mdx mice. These results are in accordance with our previously published findings in wild type mice, where we demonstrated that targeted deletion of TAK1 causes muscle atrophy *(33, 34)* whereas forced activation of TAK1 promotes muscle growth potentially through stimulating rate of protein synthesis and improving integrity of NMJs *(35)*.

A recent report showed that TAK1 is activated in skeletal muscle of DMD patients and in dystrophic muscle of mdx mice. Moreover, authors reported that inhibition of TAK1 ameliorates dystrophic phenotype in mdx mice *(36)*. However, in contrast to the published report, we found that muscle-specific inactivation/deletion of TAK1 reduces grip strength and muscle contractile function in mdx mice (**Fig. 2 and 6**). Although the exact reasons for this discrepancy in muscle phenotype remain unknown, it might be attributed to the different approaches that were used to inhibit TAK1 activity. While published report shows a small reduction in TAK1 levels after injection of AAV-TAK1 shRNA (27), our genetic approach ensures drastic reduction in the levels of TAK1 protein in dystrophic muscle of mdx mice (**Fig. 4A, 7E**). Additionally, the report shows that pharmacological inhibition of TAK1 or its knockdown using shRNA ameliorate fibrosis in mdx mice *(36)*. Contrarily, we have found that myofiber-specific ablation of TAK1 does not affect fibrosis in mdx mice (**Supplementary Fig. S3D, E**). Moreover, unlike in DMD patients where necrotic myofibers are replaced by non-contractile fibrotic tissue, in the mdx model, dramatic regenerative myogenesis program is activated and fibrosis is a secondary effect observed in diaphragm of older mdx mice *(61)*. We speculate that pharmacological or shRNA mediated approach can delete TAK1 in multiple off-target cell types which can lead to the discrepant phenotype. Indeed, a recent report shows that TAK1 controls the fibrogenic differentiation of fibro/adipogenic progenitor (FAP) cells in skeletal muscle and pharmacological inhibition of TAK1 by 5-Z-7 Oxozeaenol drastically reduced expression of fibrotic genes such as Collagen 1a1 (Col1a1), connective tissue growth factor (CTGF), periostin (Postn), Smooth muscle actin (Acta2), and fibronectin (Fn1) in FAPs *(62)*.

Even though overall muscle mass and function were diminished, we observed muscle-specific inactivation of TAK1 in young mdx mice inhibits myofiber injury evidenced by reduced area under necrosis, serum CK levels, and percentage of centronucleated myofibers (**Fig. 3**). This temporal increase in TAK1 levels during necrotic phase (∼4 week) in mdx mice is accompanied with a pronounced increase in RIPK1 phosphorylation at S321 residue (**Fig. 1E**), which is known to induce necroptosis and tissue damage *(38)*. While destabilization of sarcolemma is the major mechanism for myofiber injury, a recent report has shown that necroptosis pathway remains highly activated in the skeletal muscle of mdx mice and inhibition of necroptosis alleviates muscle damage *(39)*. Our experiments demonstrate inactivation of TAK1 drastically reduces the levels of p-RIPK1 in dystrophic muscle of 5- and 8-week-old mdx mice (**Fig. 4A**). These results indicate that in addition to reducing inflammation, inhibition of TAK1 may lessen muscle injury in young mdx mice through inhibiting myofiber necroptosis.

Autophagy is an important physiological mechanism through which defunct organelles are cleared in mammalian cells *(63)*. Targeted inhibition of autophagy causes severe myopathy in wild type mice *(64, 65)*. Interestingly, autophagy is repressed in skeletal muscle of mdx mice and in human DMD patients *(48)*. Indeed, recent reports have shown that restoring autophagy and mitophagy using pharmacological activators improves dystrophic phenotype in preclinical animal models of DMD, including mdx mice *(10, 66)*. Our results demonstrate that inducible inactivation of TAK1 in skeletal muscle of mdx mice augments the expression of components of autophagy and mitophagy (**Fig. 5**) which might be another mechanism for reduced muscle injury and improvement in muscle structure in young mdx mice.

Temporal activation of Notch and Wnt signaling is essential for regeneration of skeletal muscle upon injury *(54-56)*. Interestingly, our results demonstrate that TAK1 inactivation significantly improves gene expression of various components of Notch and Wnt signaling in dystrophic muscle of mdx mice (**Supplementary Fig. 2**). Notch signaling is essential for maintaining the quiescent satellite cell population as well as their self-renewal and proliferation *(54, 67, 68)*. In contrast, the Wnt signaling promotes differentiation of mesenchymal stem cells and satellite cells into myogenic lineage *(55, 56)*. Even though we find increased activation of Notch and Wnt signaling, there was no significant difference in PAX7 levels in dystrophic muscle of mdx;Tak1^fl/fl^ and mdx;Tak1^mKO^ mice (**Supplementary Fig.1A-C)**. By contrast, we found that inducible inactivation of TAK1 increases the expression of a few regulators of myogenic differentiation, such as MyoD and Myogenin in mdx mice (**Supplementary Fig.1G**). While the physiological significance of activation of Notch and Wnt signaling in dystrophic muscle of mdx;Tak1^mKO^ mice remain unknown, it is notable that we measured expression of various components of Wnt and Notch signaling in whole muscle, which contains multiple cell types. Therefore, it is possible that increased expression of components of Notch and Wnt signaling is a result of their activation in other cell types as well.

An important observation of the present study is that forced activation of TAK1 improves muscle growth and dystrophic phenotype in mdx mice after initial wave of myofiber necrosis. This is evidenced by our finding that overexpression of TAK/TAB1 stimulates myofiber hypertrophy in 12-week-old mdx mice. Moreover, forced activation of TAK1 also improves myofiber size and normalize fiber size distribution in young mdx mice. Similar to skeletal muscle of wild-type mice *(35)*, our experiments suggest that forced activation of TAK1 promotes muscle growth through stimulating protein translation machinery in mdx mice.

In summary, our present study demonstrates that temporal regulation of TAK1 influences disease progression in mdx mice. While more investigations are needed, our experiments provide initial evidence that activation of TAK1 supports myofiber growth in mdx mice and regulating TAK1 can be an important therapeutic approach to ameliorate disease progression and improve life span of DMD patients. Our studies are limited to one mouse model of DMD. Before trying similar strategies in clinics, more investigations are needed in higher animals, such as Golden retriever muscular dystrophy model to test whether forced activation of TAK1 ameliorates disease severity.

## MATERIAL AND METHODS

### Animals

Skeletal muscle-specific tamoxifen-inducible Tak1-knockout mice (i.e. Tak1^mKO^) were generated by crossing HSA-MCM mice (The Jackson Laboratory, Tg[ACTA1-cre/Esr1*]2Kesr/J) with floxed Tak1 (i.e. Tak1^fl/fl^) mice as described *(27, 33)*. Tak1^mKO^ mice were crossed with mdx mice to generate littermate mdx;Tak1^fl/fl^ and mdx;Tak1^mKO^ mice. Genotype of mice was determined by performing PCR from tail DNA using AccuStart II PCR genotyping kit (Quantabio, Beverly, MA). The mice were housed in a 12-h light-dark cycle and given water and food ad libitum. Inactivation of TAK1 kinase through Cre-mediated recombination in mdx;Tak1^mKO^ mice was accomplished by daily i.p. injections of tamoxifen (75 mg/kg body weight) for 4 consecutive days. After a washout period of 4 days, the mice were fed a tamoxifen-containing chow (250 mg/kg; Harlan Laboratories, Madison, WI, USA) for the entire duration of the experiment. Littermate mdx;Tak1^fl/fl^ mice (controls) were also treated with tamoxifen and fed tamoxifen-containing chow. For intramuscular delivery of adeno-associated virus (AAV), mice were anaesthetized with isoflurane and 2.5 × 10^10^ AAV vector genomes (in 30μl phosphate buffered saline, PBS) were injected in the TA muscle of mice similar to as described *(35)*. All the animals were handled according to approved institutional animal care and use committee (IACUC) protocols (protocol # 201900043) of the University of Houston.

### AAV vectors

AAVs (serotype 6) were custom generated by Vector Biolabs (Malvern, PA, USA). The AAVs expressed either GFP, Tak1 (ref seq #BC006665) or Tab1 (ref seq# BC054369) genes under a ubiquitous CMV promoter.

### Grip strength measurements

To measure total four-limb grip strength of mice, a digital grip-strength meter (Columbus Instruments, Columbus, OH, USA) was used. Before performing test, mice were acclimatized for 5 min. The mouse was allowed to grab the metal pull bar with all four paws. Tail of the mouse was then gently pulled backward in the horizontal plane until it could no longer grasp the bar. The force produced at the time of release was recorded as the peak tension. Each mouse was tested five times with a 20-40 sec break between tests. The average peak tension from five best attempts and maximum peak tension normalized against total body weight was defined as average grip strength.

### *In vivo* muscle functional assay

*In vivo* force measurements of the posterior lower leg muscles were conducted using the 1300A 3-in-1 Whole Animal System (Aurora Scientific, Aurora, Ontario, Canada) to evoke plantarflexion as described *(40)*. In brief, the mice were anaesthetized with isoflurane and placed on the isothermal stage. Optimal muscle length (Lo) that allows maximal isometric twitch force (Pt) was determined by a sequence of twitch contractions at 150 Hz every one-second with small varying changes to muscle tension and current. After muscle optimization, an average specific twitch force (sPt) was generated from 5 stimulations at 150 Hz. To obtain maximum isometric tetanic force (Po), muscles were stimulated from 25-300 Hz in 25 Hz intervals resulting in 12 specific tetanic forces (sPo). There was a 30-second delay between the first two recordings and a one-minute delay between subsequent recordings to allow sufficient muscle recovery. Following the 300 Hz stimulation, a baseline recording was obtained. Then the muscles were fatigued at 150 Hz with one contraction per second for 180 seconds. Force recordings for obtained specific isometric twitch force sPt (mN/m) and tetanic force sPo (mN/m) were normalized by total mouse body weight. For all experiments, a pre-stimulus and post-stimulus baseline of 200 ms was recorded to establish a baseline recording. For all experiments, a 0.2 ms pulse width was used. Current was adjusted on an individual basis to evoke the maximum amount of force. Contractile events were recorded using the ASI 611A Dynamic Muscle Control software (Aurora Scientific) at a sampling rate of 2000 Hz. Force (mN/m) and corresponding integral values for specific twitch force (mN/m/s) and tetanic force (mN/m/s) were calculated using the accompanying ASI 611A Dynamic Muscle Analysis software (Aurora Scientific).

### Creatine kinase (CK) assay

Levels of CK in serum were determined using a commercially available kit (Stanbio Laboratory, TX, USA) following a protocol suggested by the manufacturer.

### Histology and morphometric analysis

We performed Hematoxylin and Eosin (H&E) staining or Sirius red staining protocol on transverse diaphragm, TA, and soleus muscle sections to visualize muscle structure or fibrosis, respectively. In brief, individual hind limb muscle was isolated, flash frozen in liquid nitrogen and sectioned using a microtome cryostat. For the assessment of gross morphology, 8-10 μm thick transverse muscle sections were stained with H&E dye. Stained sections were visualized and captured using an inverted microscope (Nikon Eclipse Ti-2e microscope (Nikon), Melville, NY, USA) and a digital camera (Digital Sight DS-FI3; Nikon) at room temperature. Finally, images of the muscle sections were used for quantification of number of centronucleated myofibers (CNF), total number of myofibers and average CSA. Necrotic area in H&E-stained sections was determined by measuring percentage area filled with cellular infiltrate in the whole muscle section. The extent of fibrosis in muscle transverse sections was determined using a Picro-Sirius Red staining kit following a protocol suggested by manufacturer (Statlab, McKinney, TX). Morphometric analyses were quantified using Fiji software (U.S. National Institutes of Health, Bethesda, MD, USA).

### Immunohistochemistry

For immunohistochemistry studies, muscle sections were blocked in 1% bovine serum albumin in phosphate-buffered saline (PBS) for 1 h and incubated with primary antibodies in blocking solution at 4°C overnight under humidified conditions. For F4/80 staining, the sections were incubated with anti-F4/80 Monoclonal Antibody (BM8), PerCP-Cyanine5.5, in blocking buffer for 2h at room temperature. The sections were washed briefly with PBS before incubation with secondary Alexa tagged secondary antibody for 1 h at RT and then washed three times for 5 min with PBS. Next, the sections were stained with DAPI and the slides were mounted using a non-fluorescing aqueous mounting medium (Vector Laboratories) and and images were captured using Nikon Eclipse Ti-2e microscope (Nikon) attached with a Prime BSI Express Scientific CMOS (sCMOS) camera and Nikon NIS Elements AR software (Nikon). Damaged/permeabilized fibers in muscle cryosections were identified by immunostaining with Alexa488-or Alexa568-conjugated anti-mouse IgG. Images were stored as TIFF files and contrast levels were equally adjusted using Adobe Photoshop CS6 software (Adobe, San Jose, CA, USA). The antibodies used are provided in Supplementary Table 1.

### Western Blot

Relative levels of various proteins in skeletal muscle tissue of mdx mice was done by Western blot as previously described [32]. Briefly, skeletal muscle tissues were washed with phosphate-buffered saline (PBS) and homogenized in ice-cold lysis buffer consisting of the following: 50 mM Tris-Cl (pH 8.0), 200 mM NaCl, 50 mM NaF, 1 mM DTT, 1 mM sodium orthovanadate, 0.3% IGEPAL, and protease inhibitors. Approximately 100 μg protein was resolved in each lane on 10% to 12% SDS-polyacrylamide gels, electrotransferred onto nitrocellulose membranes, and probed using primary antibody. Signal detection was performed by an enhanced chemiluminescence detection reagent (Bio-Rad, Hercules, CA, USA). Approximate molecular masses were determined by comparison with the migration of pre-stained protein standards (Bio-Rad). Quantitative estimation of the bands’ intensity was performed using ImageJ software (U.S. NIH). The antibodies used are provided in Supplementary Table 1. The images of uncropped gel are presented in **Supplementary Fig. S6**.

### RNA isolation and qRT-PCR assay

Total RNA isolation from muscle tissues and qRT-PCR was performed as previously described *(69, 70)*. Primers for qRT-PCR were designed using Vector NTI software and their sequence has been described in our previous publications *(71-73)*.

### Statistical analyses and general experimental design

We calculated sample size using size power analysis method for a priori determination based on the s.d. and the effect size was previously obtained using the experimental procedures employed in the study. For animal studies, we estimated sample size from expected number of mdx;Tak1^mKO^ mice and mdx;Tak1^fl/fl^ mice. We calculated the minimal sample size for each group as eight animals. For some experiments, three to four animals were found sufficient to obtain statistical differences. Littermate animals with same sex and same age were employed to minimize physiological variability and to reduce SEM. The exclusion criteria for animals were established in consultation with IACUC members and experimental outcomes. In case of death, skin injury, sickness or weight loss of >10%, the mouse was excluded from analysis. Muscle tissue samples were not used for analysis in cases such as freeze artefacts on histological section or failure in extraction of RNA or protein of suitable quality and quantity. Animals from different breeding cages were included by random allocation to the different experimental groups. Animal experiments were blinded using number codes until the final data analyses were performed. Results are expressed as mean ±SEM. Statistical analyses used two-tailed Student’s t-test or one-way or two-way ANOVA followed by Tukey’s multiple comparison test. A value of p<0.05 was considered statistically significant.

## List of Supplementary Materials

Fig. S1-S6 contains supplementary figures cited in the main body of the manuscript. Table S1 contains list of the antibodies used in the study

## Acknowledgments

We thank S. Akira (Osaka University, Osaka, Japan) for providing floxed TAK1 mice.

## Author contributions

Conceptualization: AR, MW, AK

Methodology: AR, TEK, ASJ, MTS, KM

Investigation: AR, TEK, ASJ, MTS, KM

Visualization: AR, TEK, MTS

Funding acquisition: AK

Project administration: AR

Supervision: AK

Writing – original draft: AR

Writing – review & editing: AR, ASJ, MTS, MW, AK

## Data and materials availability

All data are available in the main text or the supplementary materials.

