## Supplemental Figures 1-6 and Supplemental Table 1 for "Temporal regulation of TAK1 to counteract muscular dystrophy"

**By**

**Anirban Roy, Tatiana Emy Koike, Aniket Sunil Joshi, Meiricris Tomaz da Silva, Kavya Mathukumalli, Mingfu Wu, and Ashok Kumar**

Department of Pharmacological and Pharmaceutical Sciences, University of Houston College of Pharmacy, Houston, TX 77204, USA

**Running title:** Role of TAK1 in dystrophic muscle

**Supplementary Figures S1-S6**
**Supplementary Table S1**

### SUPPLEMENTARY FIGURE LEGENDS

Supplementary Figure S1

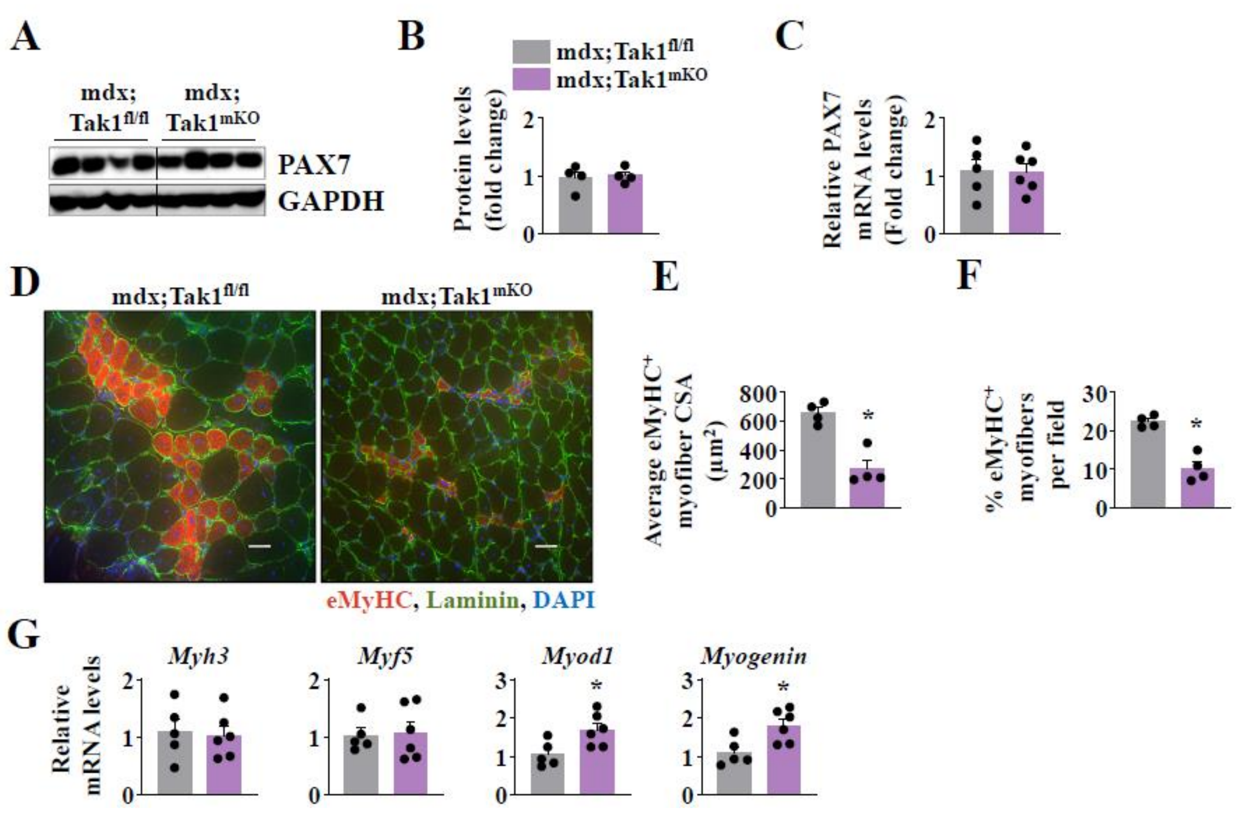


**Supplementary Fig. S1. Effect of inactivation of TAK1 on muscle regeneration in mdx mice.** 3.5 week old mdx;Tak1$^{fl/fl}$ and mdx;Tak1$^{mKO}$ mice were treated with tamoxifen and analyzed at the age of 8 weeks. (**A**) Western blot and (**B**) densitometry analysis showing Pax7 protein levels in TA muscle of 8-week old littermate mdx;Tak1$^{fl/fl}$ and mdx;Tak1$^{mKO}$ mice. (**C**) Relative mRNA levels of Pax7 in GA muscle of 8-week old mdx;Tak1$^{fl/fl}$ and mdx;Tak1$^{mKO}$ mice. Transverse sections of GA muscle was generated and immunostained for eMyHC and Laminin. Nuclei was counterstained with DAPI. (**D**) Representative photomicrographs and quantitative analysis of (**E**) average myofiber cross-section area of eMyHC$^{+}$ myofibers, and (**F**) percentage of eMyHC$^{+}$ myofibers per field in 8-week-old mdx;Tak1$^{fl/fl}$ and mdx;Tak1$^{mKO}$ mice. n=4. **(G)** Relative mRNA levels of Myh3, Myf5, Myod1, and Myogenin in skeletal muscle of 8-week-old mdx;Tak1$^{fl/fl}$ and mdx;Tak1$^{mKO}$ mice. n=5-6. Data represented as mean ± SEM. *$p \leq 0.05$, values significantly different mdx;Tak1$^{fl/fl}$ mice by Student *t* test.

Supplementary Figure S2

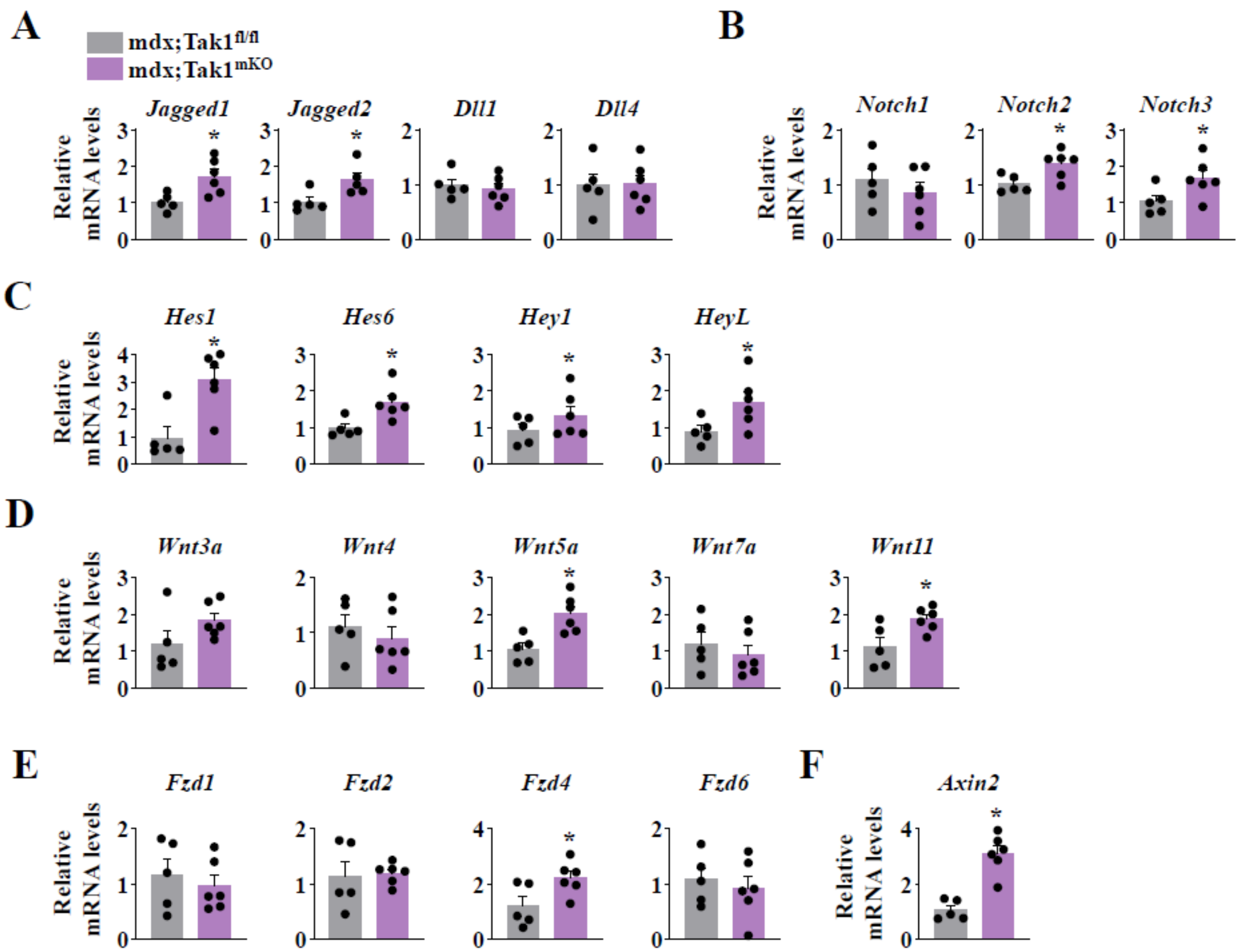


**Supplementary Fig. S2. Effect of inactivation of TAK1 on Notch and Wnt signaling in dystrophic muscle.** 3.5 week old mdx;Tak1$^{fl/fl}$ and mdx;Tak1$^{mKO}$ mice were treated with tamoxifen and analyzed at the age of 8 weeks. Relative mRNA levels of Notch ligands (**A**) Jagged1, Jagged2, Dll1, and Dll4; Notch receptors (**B**) Notch1, Notch2, and Notch3; Notch target genes (**C**) Hes1, Hes6, Hey1, and HeyL; Wnt ligands (**D**) Wnt3a, Wnt4, Wnt5a, Wnt7a, and Wnt11; Wnt receptors (**E**) Fzd1, Fzd2, Fzd4, and Fzd6; and Wnt target gene (**F**) Axin2 in GA muscle of 8-week old mdx;Tak1$^{fl/fl}$ and mdx;Tak1$^{mKO}$ mice. n=5-6. Data represented as mean ± SEM. *$p \leq 0.05$, values significantly different from GA muscle of corresponding littermate mdx;Tak1$^{fl/fl}$ mice by Student *t* test.

Supplementary Figure S3

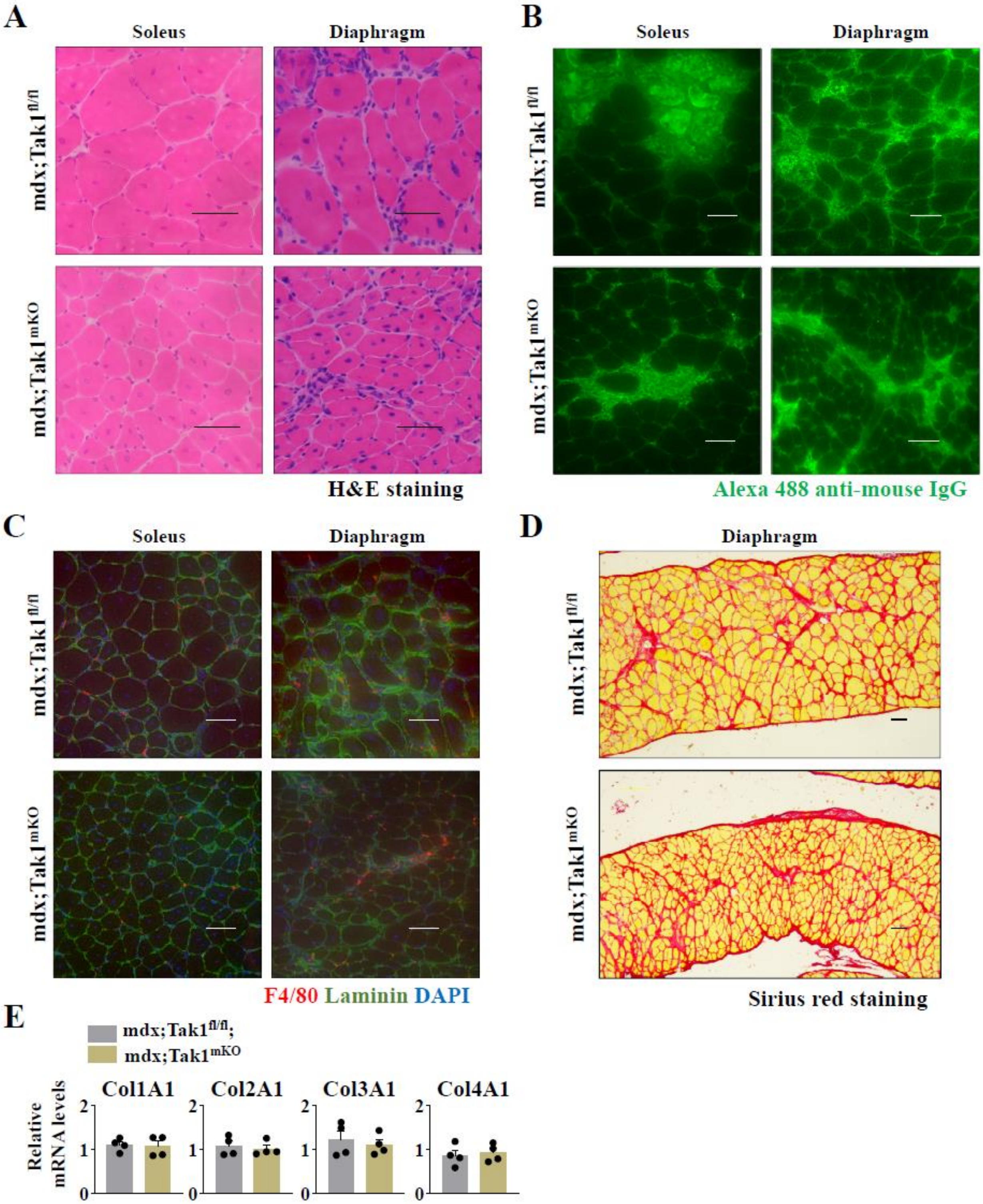


**Supplementary Fig. S3. TAK1 inactivation causes muscle wasting in adult mdx mice.** 11.5-week-old littermate mdx;Tak1$^{fl/fl}$ and mdx;Tak1$^{mKO}$ mice were treated with tamoxifen and analyzed at the age of 16 weeks. Transverse sections of soleus and diaphragm of mdx;Tak1$^{fl/fl}$ and mdx;Tak1$^{mKO}$ mice after (**A**) H&E staining, (**B**) immunostaining with Alexa488-labelled anti-mouse IgG staining, and (**C**) co-immunostaining with anti-F4/80, Laminin, and DAPI. (**D**) Transverse section of diaphragm showing Sirius Red staining in mdx;Tak1$^{fl/fl}$ and mdx;Tak1$^{mKO}$ mice. Scale bar, 50 µm. n=3. (**E**) Relative mRNA levels of *Col1A1*, *Col2A1*, *Col3A1*, and *Col4A1* in GA muscle of 16-week-old littermate mdx;Tak1$^{fl/fl}$ and mdx;Tak1$^{mKO}$ mice. n=4. Data represented as mean ± SEM. *$p \leq 0.05$, values significantly different from GA muscle of 16-week-old mdx;Tak1$^{fl/fl}$ mice by Student's t test.

**Supplementary Figure S4**

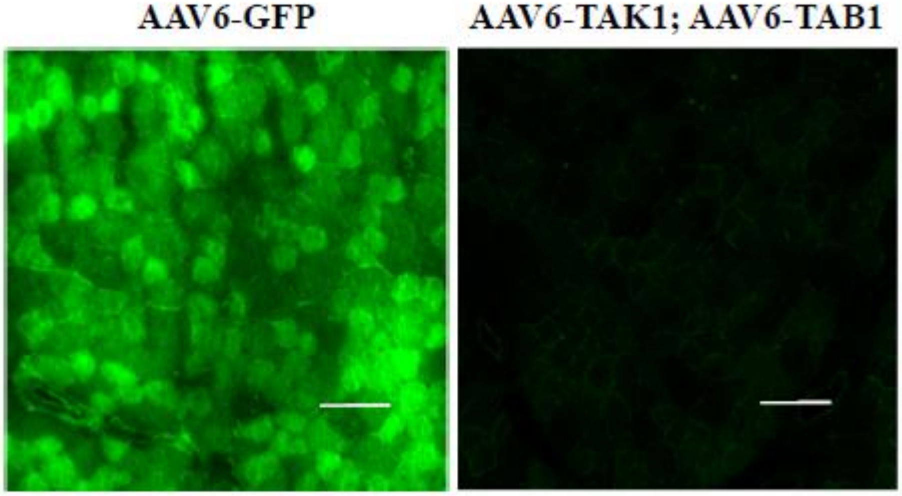


**Supplementary Fig. S4. Expression of GFP in TA muscle of mdx mice injected with AAV6-GFP.** Left side TA muscle of 12-week-old mdx mice was given intramuscular injection of AAV6-TAB1 ($1.25 \times 10^{10}$ vg) and AAV6-TAK1 ($1.25 \times 10^{10}$ vg) while the contralateral right TA muscle was injected with AAV6-GFP ($2.5 \times 10^{10}$ vg) particles. After 28 days, the mice were euthanized and the TA muscle was isolated and analyzed for GFP expression. Scale bar, 100 µm.

**Supplementary Figure S5**

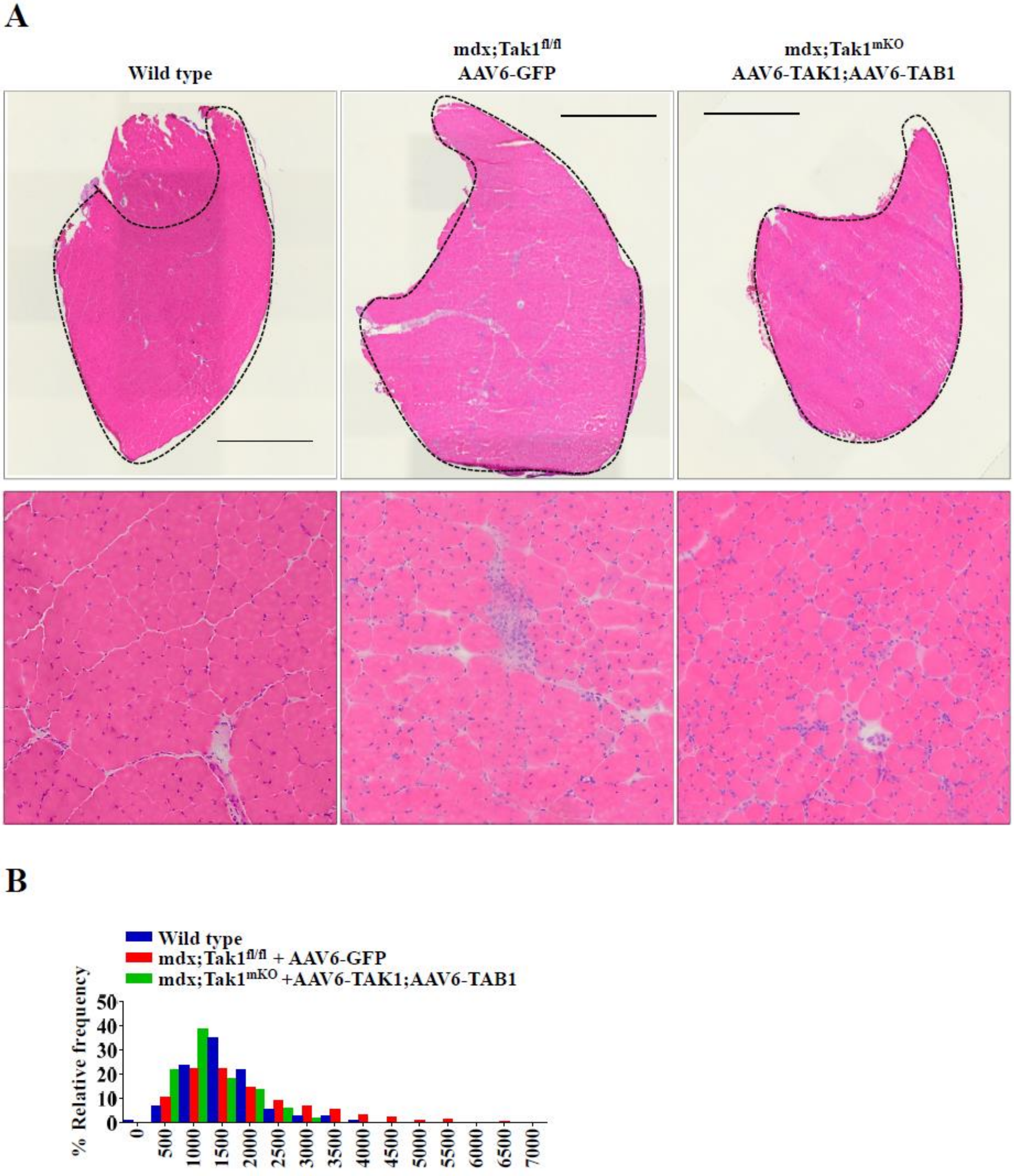


**Supplementary Fig. S5. Temporal regulation of TAK1 improves myopathy in mdx mice.** 3.5-week-old mdx;Tak1$^{fl/fl}$ and mdx;Tak1$^{mKO}$ mice were treated with tamoxifen for four days. At 6 weeks, the left side TA muscle was given intramuscular injection of AAV6-TAB1 ($1.25 \times 10^{10}$ vg) and AAV6-TAK1 ($1.25 \times 10^{10}$ vg) while the contralateral right TA muscle was injected with AAV6-GFP ($2.5 \times 10^{10}$ vg) particles. At the age of 10 weeks, the mice were euthanized and the

TA muscles were harvested and analyzed. (**A**) H&E-stained transverse sections of whole TA muscle (top panel) and magnified inset (bottom panel) of age matched wt mice, mdx;Tak1$^{fl/fl}$ mice injected with AAV6-GFP, and mdx;Tak1$^{mKO}$ mice co-injected with AAV6-TAK1 and AA6-TAB1. (**B**) Relative frequency distribution showing myofiber CSA of wild type, mdx;Tak1$^{fl/fl}$ mice injected with AAV6-GFP, and mdx;Tak1$^{mKO}$ mice co-injected with AAV6-TAK1 and AAV6-TAB1.

Supplementary Figure S6

Fig. 1A

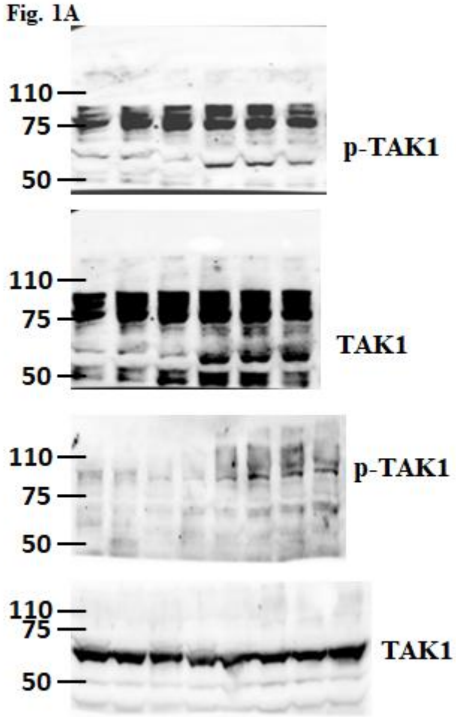


Fig. 1E

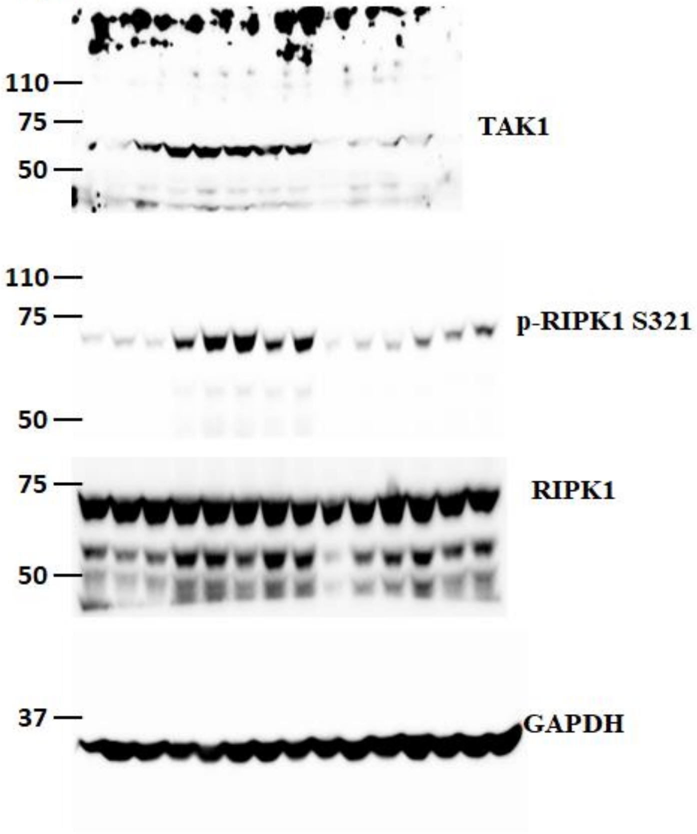


Fig. 4A

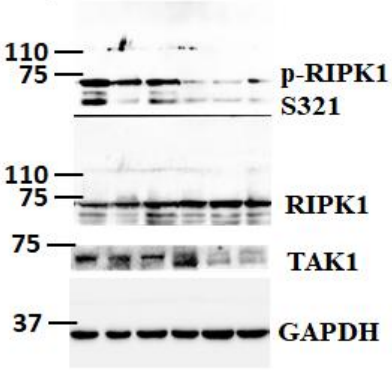


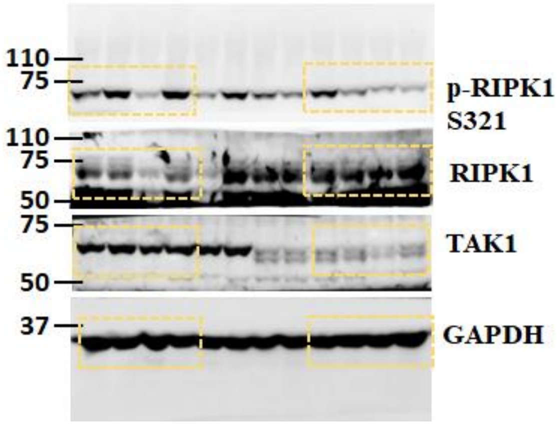

Supplementary Figure S6 contd.

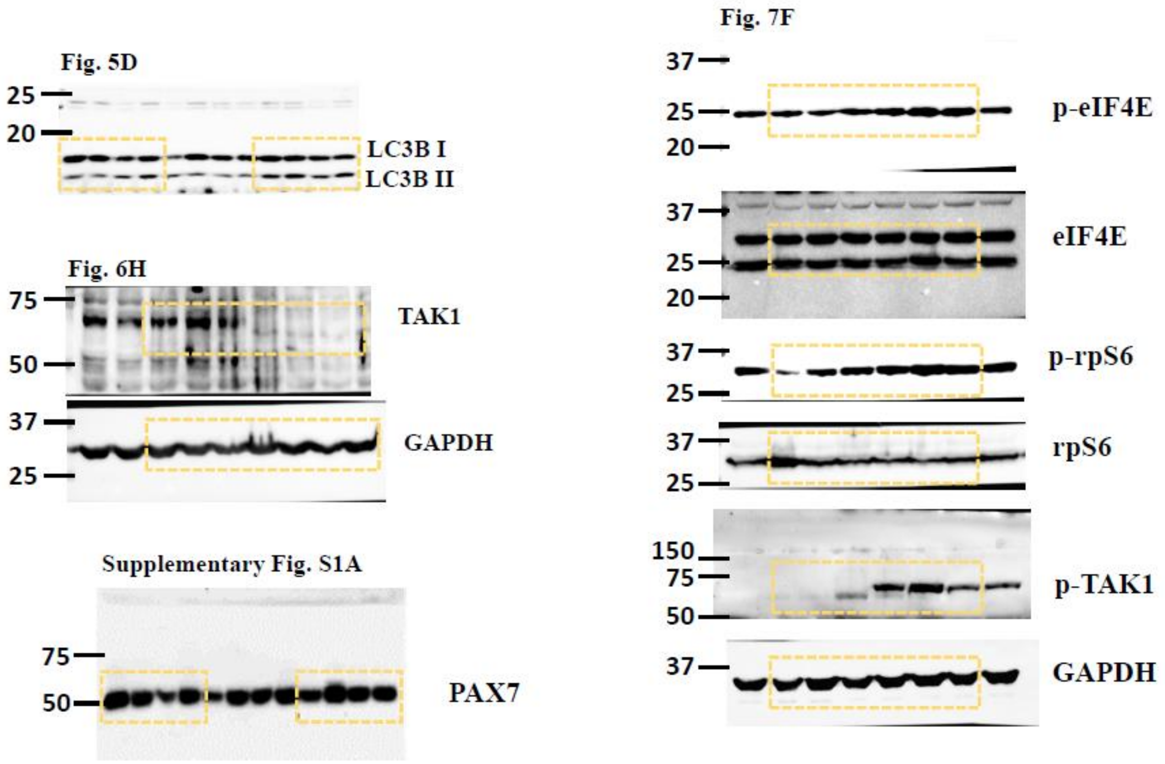


**Supplementary Fig. S6. Uncropped images of immunoblots.** Raw immunoblots and bands shown in original figures are marked here.

**Supplementary Table 1**. Antibodies used in the study.

| **Antibody** | **Source and Catalog no.** |
|---|---|
| Monoclonal rabbit-anti-total-TAK1 | Cell Signaling Technology, # 5206 |
| Monoclonal rabbit-anti-phospho-TAK1 | Invitrogen #MA5-15073 |
| Monoclonal rabbit-anti-phospho-RIP | Cell Signaling Technology, # 83613 |
| Monoclonal rabbit-anti-RIP1 | Cell Signaling Technology, # 3493 |
| Monoclonal rabbit-anti-LC3B | Cell Signaling Technology, # 3868 |
| Monoclonal rabbit-anti-GAPDH | Cell Signaling Technology # 2118 |
| Polyclonal rabbit-anti-phospho-eIF4E | Cell Signaling Technology # 9741 |
| Monoclonal rabbit-total-eIF4E | Cell Signaling Technology # 2067 |
| Monoclonal rabbit-anti-phospho-S6 Ribosomal Protein | Cell Signaling Technology # 4858 |
| Monoclonal rabbit-anti-total-S6 Ribosomal Protein | Cell Signaling Technology # 2217 |
| Polyclonal rabbit-anti-Laminin | Sigma, L9393 |
| Monoclonal mouse-anti-Pax7 | DSHB Cat# pax7 |
| Monoclonal mouse-anti-eMyHC | DSHB Cat# F1.652 |
| Monoclonal mouse-anti-F4/80 (BM8) | Invitrogen Cat# 45-4801-82 |
| Polyclonal goat-anti-rabbit IgG Alexa Fluor 568 | Invitrogen # A-11036 |
| Polyclonal goat-anti-mouse IgG Alexa Fluor 568 | Invitrogen # A-11004 |
| Polyclonal goat-anti-mouse IgG Alexa Fluor 488 | Invitrogen # A32731 |
| Polyclonal goat-anti-mouse IgG Alexa Fluor 555 | Invitrogen # A-21127 |
| Polyclonal goat-anti-rabbit IgG Alexa Fluor 488 | Invitrogen # A-11034 |
